# Convergence and horizontal gene transfer drive the evolution of anaerobic methanotrophy in archaea

**DOI:** 10.1101/2024.05.23.595608

**Authors:** Philip H. Woods, Daan R. Speth, Rafael Laso-Pérez, Daniel R. Utter, S. Emil Ruff, Victoria J. Orphan

**Author notes:** Division of Microbial Ecology, Centre for Microbiology and Environmental Systems Science, University of Vienna, Vienna, Austria (Daan R. Speth).

## Abstract

Despite their large environmental impact and multiple independent emergences, the processes leading to the evolution of anaerobic methanotrophic archaea (ANME) remain unclear. This work uses comparative metagenomics of a recently evolved but understudied ANME group, ‘*Candidatus* Methanovorans’ (ANME-3) to identify evolutionary processes and innovations at work in ANME which may be obscured in earlier evolved lineages. Within members of *Methanovorans*, we identified convergent evolution in carbon and energy metabolic genes as likely drivers of ANME evolution. We also identified erosion of genes required for methylotrophic methanogenesis along with horizontal acquisition of multi-heme cytochromes and other loci uniquely associated with ANME. The assembly and comparative analysis of multiple Methanovorans genomes offers important functional context for understanding the niche-defining metabolic differences between methane-oxidizing ANME and their methanogen relatives. Furthermore, this work illustrates the multiple evolutionary modes at play in the transition to a novel and globally important metabolic niche.

## 2 Introduction

Anaerobic methanotrophic archaea (ANME), originally identified in 1999 (*1*, *2*, *3*, *4*), are now known to be globally distributed (*5*) and play a major role in marine cycling of the potent greenhouse gas methane (*6*). Marine environments account for up to 25% of global annual methane production, but ANME act as a control on marine methane flux by consuming 30% of global production before it reaches the water column (*7*). In marine ecosystems, ANME perform anaerobic oxidation of methane (AOM) coupled to sulfate reduction in syntrophy with several recently defined clades of sulfate-reducing bacteria (SRB) (*2*, *4*, *8*, *9*). Despite their global environmental impact and several insights into ANME metabolism, important details regarding their physiology, ecology, and evolution remain incompletely understood (*10*, *11*, *12*, *13*, *14*, *15*, *16*, *17*).

Phylogenetic, genomic, and physiological similarities between methanotrophic ANME and methanogenic archaea within the *Methanosarcinaceae* have led to the hypothesis that ANME evolved from methanogens. Though they do not form a singular ANME clade, ‘*Candidatus* Methanovorans’ (ANME-3), ‘*Candidatus* Methanocomedenaceae’ (ANME-2ab), and ‘*Candidatus* Methanogasteraceae’ (ANME-2c) are closely related to methanogens in 16S rRNA gene and concatenated marker gene phylogenies, and have many central metabolic pathways and energy-conserving complexes in common with methanogens (*10*, *11*, *12*, *18*, *14*, *16*). For example, ANME accomplish AOM by ‘reverse methanogenesis’, using the Mcr enzyme, responsible for the final step in methanogenesis, to initiate methane oxidation (*10*, *11*, *19*). Also, ANME-2, ANME-3, and cytochrome-containing methanogens are predicted to use the same electron carriers, such as the membrane-bound molecule methanophenazine, though this has not yet been shown definitively (*16*). In addition to these biochemical similarities, some methanogens may be able to oxidize methane under certain conditions (*20*, *21*, *22*, *23*), and minor back flux of carbon during AOM has been reported under natural conditions (*24*). Though there is not yet strong evidence that backwards carbon flow can be coupled to growth in either methanogens or ANME in their natural state, these observations indicate that the enzymes of the canonical methanogenesis/AOM pathway are reversible. While these observations strongly suggest ANME have methanogenic ancestors, they do not indicate how such an evolutionary transition may have occurred.

Though ANME are polyphyletic, suggesting that each clade may have evolved independently, there may be commonalities in the processes and pressures leading to each clade’s evolution. Importantly, phylogenetically distinct ANME clades share many common features not present in methanogens. These include the apparent acquisition of large multiheme cytochrome c proteins hypothesized to be associated with extracellular electron transfer (EET), and soluble protein complexes possibly used for electron transfer and recycling of electron carriers (*16*). However, it has been difficult to identify the evolutionary processes involved in ANME evolution (*16*). Investigations have been impeded by a lack of sufficient high-quality genomes and the long divergence times between previously studied ANME and their closest methanogen relatives. These previously studied ANME include the *Methanophagales* (ANME-1), whose nearest methanogen relatives are in a separate taxonomic order, and several subtypes of ANME-2, whose nearest methanogen relatives are in separate taxonomic families. While these long divergence times could mean that these ANME genomes are highly optimized for methanotrophy through loss of extraneous genes, they also complicate attempts to infer the last common ancestral state between these ANME and methanogens. For example, long divergence times between organisms typically allows evolutionary “noise” to accumulate in their genomes. The combination of these factors has made it nearly impossible to identify a minimal set of changes required to shift from methanogenic to methanotrophic lifestyles using genomes of earlier evolved Methanocomedens (ANME-2a), Methanomarinus (ANME-2b), and Methanogaster (ANME-2c).

In contrast, study of the recently described *Methanovorans* (ANME-3) clade may provide insights necessary to clarify the process of ANME evolution. Based on 16S rRNA gene phylogeny, *Methanovorans* cluster as a genus within the family *Methanosarcinaceae*, implying they may have emerged more recently than other ANME groups (*25*). A recent study of the first two assembled *Methanovorans* genomes supported this observation, but was unable to draw conclusions about *Methanovorans* evolution due to the small sample size (*16*). In the present study, we compare ten *Methanovorans* (ANME-3) genomes from three sites with representative genomes of *Methanocomedens*, *Methanomarinus*, and *Methanogaster* (ANME-2a, -2b, and -2c) and methanogenic *Methanosarcinaceae* (Tab. S1). These *Methanovorans* genomes include *de novo* metagenome-assembled genomes (MAGs) and previously published (though sometimes unclassified due to missing 16S rRNA genes) *Methanovorans* MAGs (*16*, *26*). Through this comparison, we identify relevant examples of gene loss, horizontal transfer, and convergent evolution in *Methanovorans* genomes during their evolution toward methanotrophy, and present a conceptual model for ANME evolution. These findings identify key elements for future investigation of ANME physiology and provide perspective into the processes involved in microbial evolution of novel metabolisms and novel symbioses.

## 3 Results

A reference set of ten high-quality *Methanovorans* (ANME-3) MAGs (metagenome-assembled genomes) formed the backbone of this analysis, reconstructed from locations at the Haakon Mosby Mud Volcano in the Barents Sea, Jaco Scar on the Costa Rica Pacific margin, and the Scotian Basin in the North Atlantic. These *Methanovorans* MAGs are the largest collection of genome-scale data on the genus to date, and are derived from published data (*16*), re-binning of previously misidentified genomes (*26*), and *de novo* assembly of metagenome data obtained for this study. The collection of genomes has a median completeness and redundancy of 84.45% and 1.5% respectively, and average nucleotide identity calculations suggest these genomes represent five distinct species (ANI of 81.2%–99.8%) (Fig. 1, Tab. S1). Three of these species are represented by 2–4 genomes (Tab. S1). Though two of the species are represented by only a single genome, these genomes (MAGs HMMV and HMMV2) have high estimated quality (Tab. S1). Interestingly, the species divisions within *Methanovorans* do not reflect the sampling site geography. For example, *Methanovorans* sp. HMMV1 (from the Haakon Mosby Mud Volcano in the Barents Sea) is more closely related to species from Jaco Scar (on the Costa Rica Pacific margin) than to the other Haakon Mosby Mud Volcano species (*Methanovorans* sp. HMMV2).

**Figure 1.**
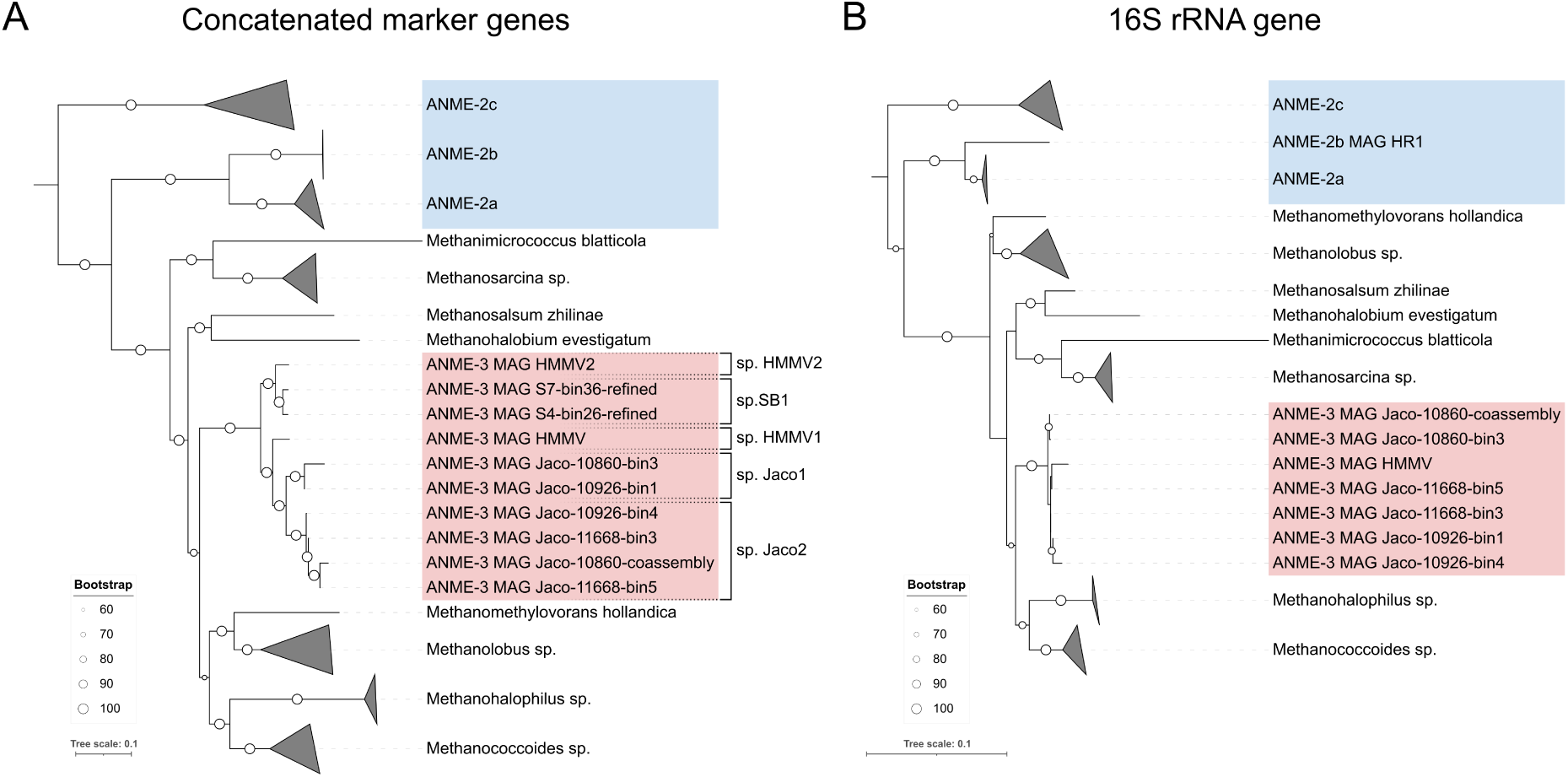
Estimated species phylogeny resulting from (A) the concatenated alignment of 60 archaeal marker genes and (B) the 16S rRNA gene. The root of both trees leads to *Methermicoccus shengliensis* DSM 18856. *Methanocomedenaceae* (ANME-2ab) and *Methanogasteraceae* (ANME-2c), highlighted blue, sit outside of the *Methanosarcinaceae*, while *Methanovorans* (ANME-3), highlighted red, sit among them. The placement of *Methanovorans* within this family is slightly different in each tree, but the 16S tree has low bootstrap support for several internal nodes of the *Methanosarcinaceae*. The ten *Methanovorans* MAGs can be grouped into five distinct species based on average nucleotide identity (*>* 95% ANI) which are well recovered by the concatenated marker gene tree, indicated by black brackets.

As observed in previous studies, we find the *Methanovorans* (ANME-3) are closely related to methanogens, clustering within the widespread and metabolically versatile family *Methanosarcinaceae* (Fig. 1) (*25*, *16*). While the association of *Methanovorans* with the *Methanosarcinaceae* is robust, their exact position within this methanogenic family appears to vary depending on the gene(s) used to construct the phylogeny. 16S rRNA gene trees tend to place their divergence closer to the tips of the phylogeny, while concatenated gene trees tend to place them closer to the base of the *Methanosarcinaceae* (Fig. 1) (*25*, *16*). Unlike the 16S rRNA gene phylogeny, the concatenated marker gene phylogeny of *Methanovorans* independently recovers the species clusters identified through ANI analysis (Fig. 1). In either case, our expanded set of *Methanovorans* genomes confirms that they are the ANME group with the most recent common ancestor with methanogens. *Methanovorans* genomes encode genes which are significantly overrepresented relative to related methanogens in the COG categories mobile elements (X) and defense mechanisms (V), as well as multi-heme cytochrome (MHC) proteins (supp. text, file S1). *Methanovorans* and other ANME also appear to share many distinctive transporter proteins either enriched relative to or not found in related methanogens, including a highly conserved putative lipoprotein export system (supp. text). Detailed investigations of individual protein complexes indicated that large ANME MHCs, contractile injection systems, nitrogenase, and phosphonate-metabolizing C-P lyase were horizontally transferred into *Methanovorans*, whereas key central metabolic pathways including the methane-metabolizing complex Mcr (methyl-coenzyme M reductase) and the energy-conserving respiratory complex Rnf (Rhodobacter nitrogen fixation) exhibit signs of sequence-level convergent evolution between ANME-2abc and ANME-3 (discussed below). Several other proteins potentially involved in ANME energy metabolism (e.g. Hdr as noted by Chadwick *et al.* (*16*)) appear to be late horizontal acquisitions in *Methanovorans*. Additionally, all investigated ANME except three of the ANME-3 MAGs lack genes involved in methylotrophic methanogenesis (discussed below).

### 3.1 Formerly core pathways for methylotrophic methanogenesis have been partially lost in *Methanovorans*

Within the *Methanosarcinales*, the family *Methanosarcinaceae* is distinctive for its unique ability to perform methanogenesis from methylated amine C1 compounds (*27*). This pathway requires the enzymes MtbABC, MtmBC, MttBC, and RamA (*28*). Several of the enzymes in this pathway (MtmB, MtbB, and MttB, the methyltransferases for mono-, bi-, and trimethylamine, respectively) also require the non-canonical amino acid pyrrolysine and its associated biosynthetic genes (supp. text) (*28*). All methanogens within the *Methanosarcinaceae* family share this metabolism, so it is parsimonious to assume that the methanogenic ancestor of *Methanovorans* was capable of methylamine metabolism and pyrrolysine biosynthesis. However, we expect that these genes will be lost by *Methanovorans* in their transition to methanotrophy due to the presumed limited use for the genes in methanotrophic metabolism.

In support of this transition from methylotrophic methanogenesis to methane oxidation, three of the ten studied *Methanovorans* genomes retained relicts of the methylamine metabolism and pyrrolysine biosynthesis pathways. (Fig. 2). Two of these (sp. SB1, MAG S4-bin26-refined and MAG S7-bin36-refined) retain genes for both pathways (Fig. S1) while the third (sp. HMMV1, MAG HMMV) retained a partial pathway for pyrrolysine biosynthesis and no genes in the methylamine pathway. *Methanovorans* MAG HMMV pyrrolysine biosynthesis genes showed signs of severe degradation including insertions and premature truncation (Fig. 2 and supp. text). These features suggest that the pathway is not functional in *Methanovorans* MAG HMMV. The capacity for methanogenesis from methanol, another common metabolism in *Methanosarcinaceae*, could not be conclusively identified in any analyzed ANME genome (supp. text).

**Figure 2.**
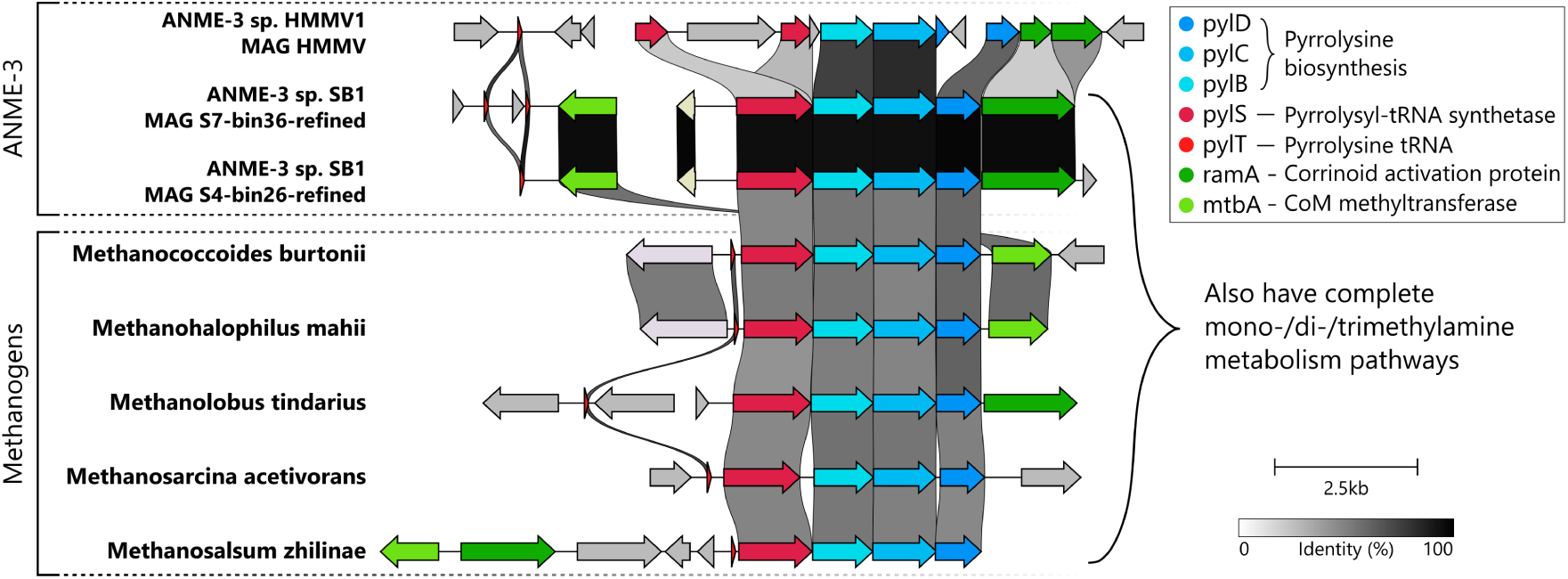
Gene synteny of pyrrolysine metabolism in *Methanovorans* (ANME-3) and representative methanogens. All methanogen genomes contain genes for pyrrolysine biosynthesis (blue) and expression (red) and methanogenesis from methylamines (green). Two *Methanovorans* genomes contain these genes, seven *Methanovorans* lack them completely, and one *Methanovorans* contains a partial set of genes with several disruptions in the gene sequences.

Codon usage in all ten *Methylovorans* genomes further supports that pyrrolysine-utilizing methylotrophic pathways have been lost or are no longer used. In methylotrophic, pyrrolysine-utilizing *Methanosarcinaceae*, pyrrolysine is encoded using the TAG stop codon (*29*). This codon is rarely used for normal termination of translation in pyrrolysine-utilizing organisms, but it is much more common in related taxa which lack pyrrolysine (*30*). The pyrrolysine-utilizing methanogens in this study used TAG for less than 5% of their stop codons on average, but the methanotrophic ANME-2 lineages *Methanocomedenaceae* and *Methanogasteraceae* have over three times the TAG stop codons (18% of their genes, much like non-pyrrolysine-utilizing methanogens) (Tab. S2). All *Methanovorans*, including the two genomes with complete methylotrophy and pyrrolysine synthesis pathways, have a TAG codon abundance of about 17%, closely matching the that of non-pyl-utilizing taxa (Tab. S2).

### 3.2 Core methanotrophy pathways in *Methanovorans* have converged with Methanocomedenaceae *or* Methanogasteraceae

ANME MCR protein sequences share conserved residues distinct from methanogens. The methyl-coenzyme M reductase (MCR) complex catalyzes the first step in the anaerobic oxidation of methane (or the last step in methanogenesis). We find that phylogenies of the three structural subunits of Mcr consistently place *Methanovorans* between other methanotrophic taxa and the methanogenic *Methanosarcinaceae*, in contrast to their position within the *Methanosarcinaceae* in consensus species trees (Fig. 3). At a glance, it is not clear what sequence features might be responsible for the closer grouping between *Methanocomedenaceae* (ANME-2ab) and *Methanogasteraceae* (ANME-2c) (ANME-2abc) and *Methanovorans* (ANME-3) MCR. However, there are individual residues throughout the protein sequences which have high levels of conservation (*>* 70% identity, *<* 30% gaps) among the studied ANME while simultaneously being different from all residues at those positions in methanogen-encoded MCRs. The rate of these conserved, ANME-characteristic residues is several times higher in MCR protein sequences (5.8 per kAA) than in the conserved phylogenetic markers used to estimate the species phylogeny (2.6 per kAA) (Tab. S3, supp. text). The location of these residues in predicted structures of MCR are generally not near the active site, making their function unclear (supp. text). Additionally, analysis of codon substitution rates using the HyPhy software’s aBSREL method to test for branch-specific selection indicate that all three structural subunits of MCR (McrABG), but not the non-structural *mcrC* and *mcrD*, experienced positive selection during the divergence of the *Methanovorans* genus (HyPhy aBSREL, *mcrA*: *p <* 0.00011, *mcrB* : *p <* 0.00018, *mcrG* : *p <* 0.019).

**Figure 3.**
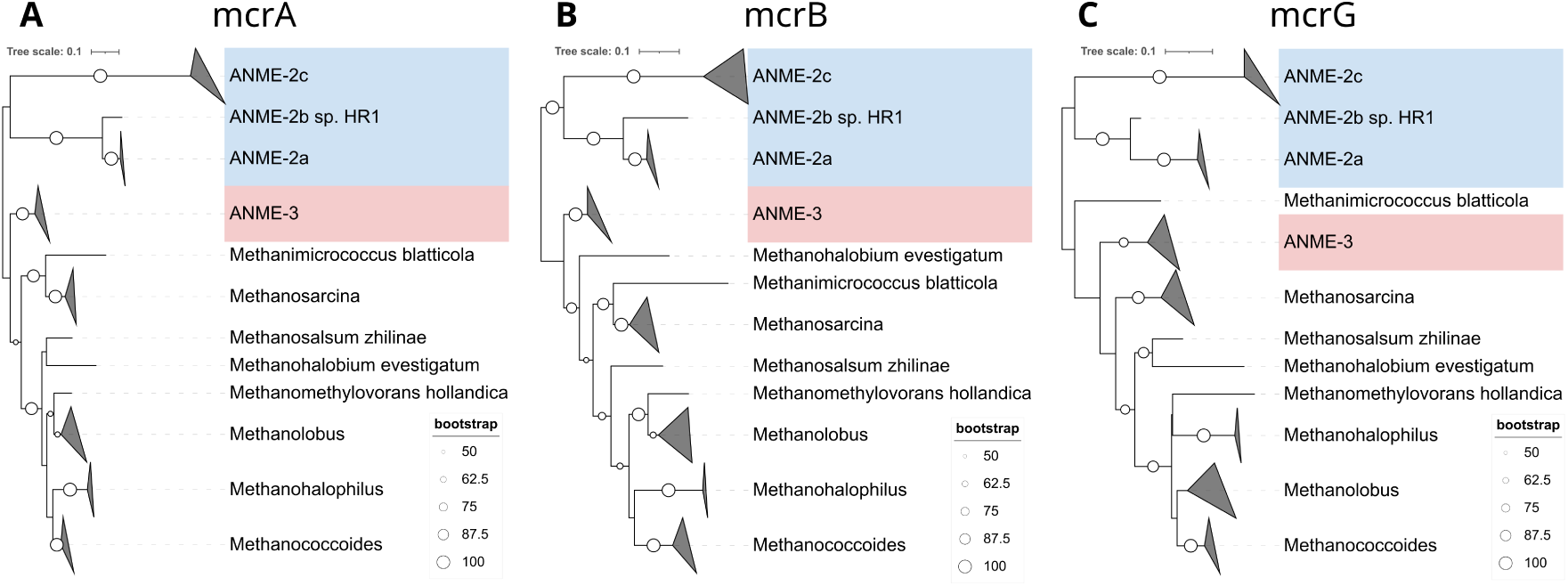
Gene phylogeny of MCR structural subunits mcrA (A), mcrB (B), and mcrG (C). Compared to the estimated species phylogeny, the *Methanovorans* (ANME-3) clade is positioned closer to *Methanocomedenaceae* (ANME-2ab) and *Methanogasteraceae* (ANME-2c) in each tree except the mcrD phylogeny (not shown). However, ANME-3 do not cluster within *Methanocomedenaceae* or *Methanogasteraceae*, and generally sit closer to methanogenic taxa than to either methanotrophic group.

ANME energy conservation genes also share distinctive sequence and structural features indicative of convergent evolution. The RNF complex is a reversible energy-conserving system found in a variety of organisms including diverse bacteria, methanogenic archaea, and ANME which couples sodium ion transport to redox transformations of ferredoxin (*31*). Though the role of this complex in ANME is unclear, its typical role in low-energy metabolism (e.g. (*32*)) and conservation in related methanogens makes it an intriguing target for investigation.

*Methanovorans* share identical gene synteny with *Methanocomedenaceae* within most of the RNF locus (Fig. 4, S2). However, *Methanovorans* retain an uncharacterized conserved protein between *apbE* and the rest of the locus otherwise present only in their methanogenic relatives. *Methanovorans* protein sequences from this locus tend to share slightly higher amino acid identity with methanogens than with *Methanocomedenaceae* or *Methanogasteraceae*. Also, the *Methanovorans* cytochrome b protein and the short hairpins have only low to moderate sequence identity to their *Methanocomedenaceae* counterparts. Surprisingly, the RNF genes do not all seem to share the same phylogeny. While phylogenies of some genes in the locus concord well with the estimated species tree, phylogenies of others indicate that *Methanovorans* fall between their methanogenic relatives and the methanotrophic *Methanocomendaceae* and *Methanogasteraceae* (Fig. 4, S3). Though *Methanovorans* (ANME-3) RNF protein sequences share higher amino acid identity with RNF in methanogens, they often exhibit localized sequence and/or structural features more similar to *Methanocomendaceae* (ANME-2ab) or *Methanogasteraceae* (ANME-2c). One such example is the N-terminal RnfC barrel sandwich hybrid domain, which appears to have different domain annotation and potentially different structural features in ANME (supp. text, Fig. S4, Pfams in file S1). Another striking example of similar domain features between ANME is the multi-heme cytochrome protein MmcA. These features include small deletions in *Methanovorans* in a similar position to *Methanocomedenaceae* sequences and a highly conserved heme binding motif which is altered from methanogens (supp. text, Fig. S5). However, the protein sequence also shares several domain features and higher overall similarity with the methanogenic *Methanosarcinaceae*, suggesting the common features between *Methanovorans* and *Methanocomedenaceae* represent convergent evolutionary modifications rather than horizontal gene transfer (supp. text, Fig. S5).

**Figure 4.**
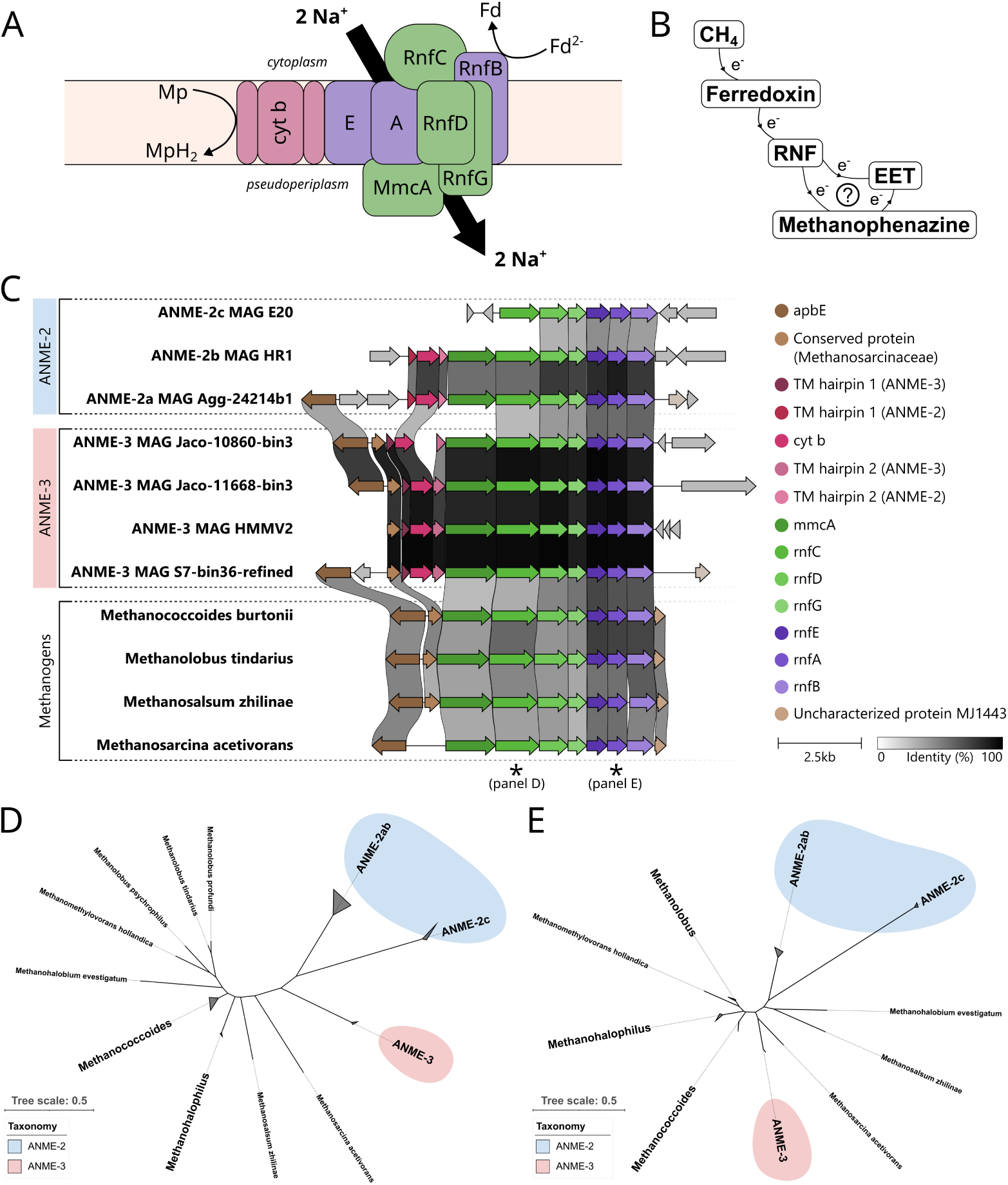
RNF complex organization (A), position in ANME metabolism (B), locus synteny (C), and gene phylogeny of *rnfC* (D) and *rnfA* (E). In panels A and C, subunits or genes colored purple concord with the estimated species phylogeny, subunits or genes colored green do not, and subunits or genes colored red are part of the locus only in ANME. The RNF complex (A) is seated within the membrane and couples electron transfer to sodium ion translocation. Gene organization of the rnf locus (C) is similar but not identical in ANME and methanogens. All studied genomes had the same organization of rnfCDGEAB and mmcA with the exception of *Methanogasteraceae* (ANME-2c). In addition, *Methanocomedenaceae* (ANME-2ab) and *Methanovorans* (ANME-3) loci also have two small transmembrane hairpin proteins and a b-type cytochrome, though the sequence similarity of these genes between *Methanovorans* and *Methanocomedenaceae* is low. *Methanovorans* and methanogenic *Methanosarcinaceae* have conserved flanking genes upstream of the locus, despite the extra genes in the *Methanovorans* loci. The gene phylogeny of rnfC (D) differs from the estimated species phylogeny, while the phylogeny of rnfA (E) matches the species phylogeny more closely.

### 3.3 Conserved modes of ANME interspecies syntrophic interaction are gained by horizontal gene transfer in *Methanovorans*

In contrast to the previously discussed core metabolism genes, other metabolic genes reflect potential horizontal gene transfer events from ANME. For example, *Methanovorans* (ANME-3) appear to have gained large MHC proteins, previously hypothesized to facilitate direct interspecies electron transfer (*33*, *34*), after their last common ancestor. Previous work has noted that ANME genomes often encode very large multiheme cytochrome c proteins (*16*) and proposed three major subclasses (MHC-A, MHC-B, and MHC-C) based on the presence and position of domains. Many *Methanocomendaceae* (ANME-2ab) and *Methanogasteraceae* (ANME-2c) genomes encode one representative of each subclass (*16*), but *Methanovorans* do not. One subclade of *Methanovorans* contains only MHC-A, while another *Methanovorans* subclade contains only MHC-B and MHC-C and/or a novel class which we are calling MHC-D (Fig. 5). The MHC-A gene in *Methanovorans* genomes has similar domain architecture to previously described MHC-A. However, it is about twice the size of the gene in other ANME except for a single *Methanocomedens* (ANME-2a) genome (MAG Agg-24214b1), and sequence similarity is relatively low between *Methanovorans* MHC-A and other ANME (supp. text). The MHC-B/MHC-C locus in *Methanovorans* has good sequence identity (MHC-B: 89%, MHC-C: 74%) and high synteny with *Methanocomedens* genomes (Fig. 5). However, MHC-D has domain architecture unlike recognized genes in other ANME (supp. text).

**Figure 5.**
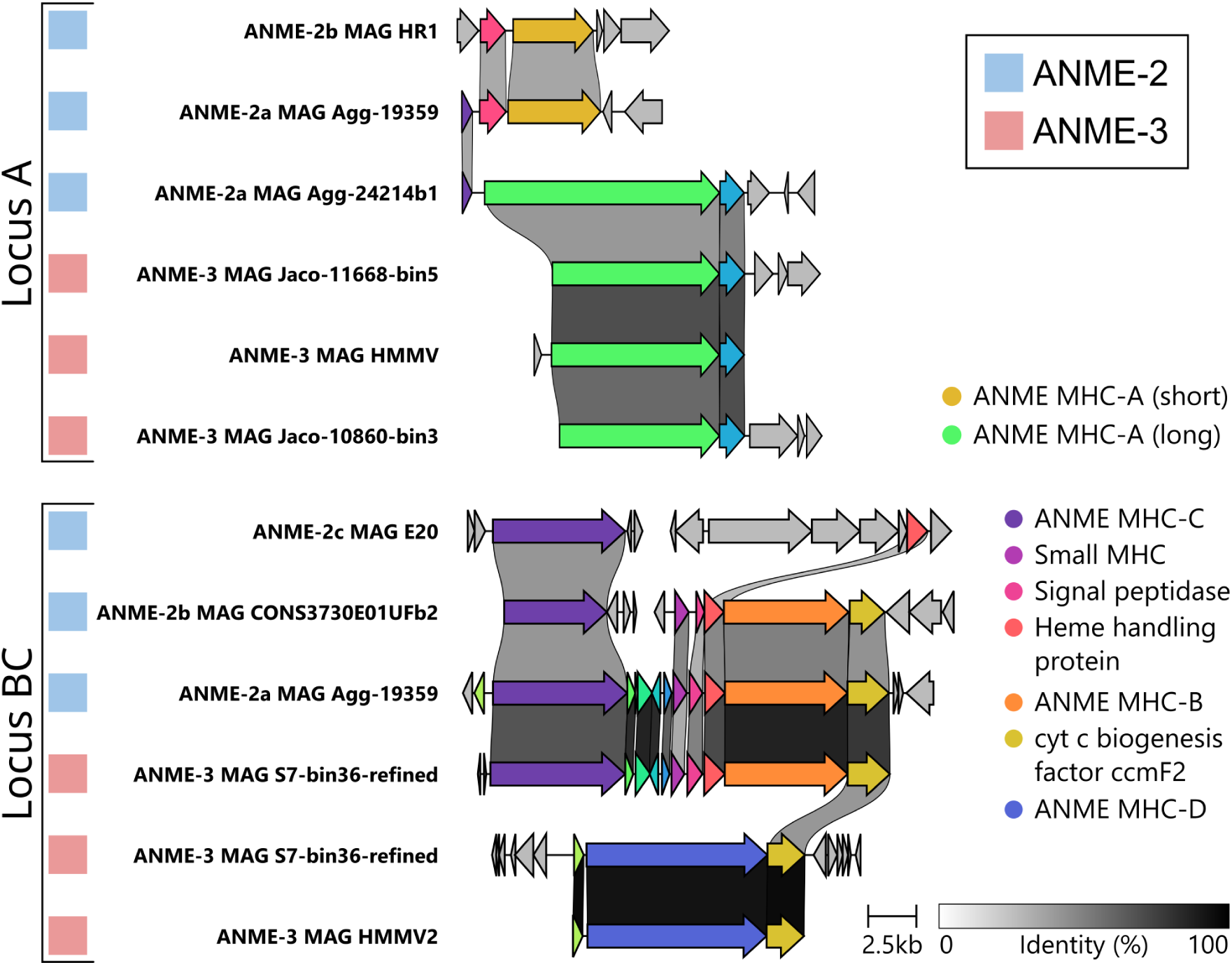
*Methanovorans* (ANME-3) contain large multi-heme cytochrome proteins (MHC-A, -B, -C) typical of other ANME clades. MHC-A and MHC-BC have distinct genomic loci in *Methanovorans* which are typically most similar to *Methanocomedens* (ANME-2a) species in sequence and organization. Each *Methanovorans* species has at least one genome representative containing large ANME MHCs, but *Methanovorans* never have both loci in the same genome. The *Methanovorans* species can be split into two monophyletic clades, one with MHC-A loci and one with MHC-BC loci. Two *Methanovorans* genomes contain a large MHC protein apparently resulting from the fusion of the N-terminus of MHC-C and the C-terminus of MHC-B.

Contractile injection systems (CIS) in *Methanovorans* also appear to be horizontally acquired. Phage-like protein translocation systems (PLTS) are a subtype of CIS which have been previously noted as a common feature of ANME genomes which may have different phylogenetic histories from the core genome (*16*). Of the ten *Methanovorans* genomes in this study, seven have PLTS components and six appear to have a complete injection system. Based on the phylogeny of several core PLTS structural genes, *Methanovorans* cluster most closely with *Methanogaster* (ANME-2c) (Fig. S6). *Methanovorans* CIS loci also appear to have higher amino acid identity with *Methanogaster* CIS than with those of other archaea (Fig. 6). However, the genes in *Methanogaster* CIS loci represented in this study are typically not more than 60% identical to the corresponding *Methanovorans* genes.

**Figure 6.**
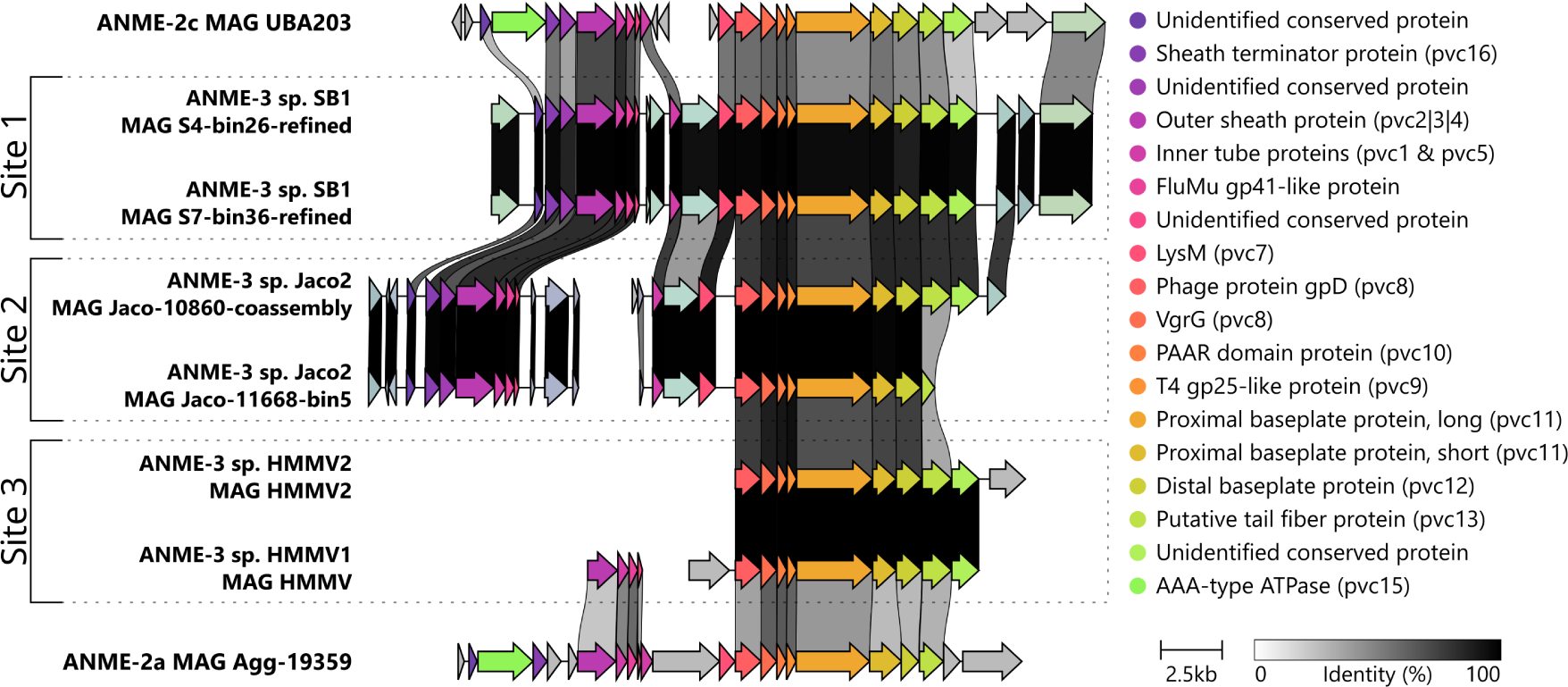
Gene synteny of contractile injection system loci in *Methanocomedens* (ANME-2a), *Methanogaster* (ANME-2c), and *Methanovorans* (ANME-3). Colors denote groups of genes with at least 30% amino acid identity. Gray links between genes are colored according to their pairwise amino acid identity. *Methanovorans* CIS loci bear most similarity in sequence identity and gene organization to CIS loci in *Methanogaster*. *Methanovorans* have three distinct subtypes of CIS loci, as determined by sequence similarity, gene content, and locus synteny. These subtypes correspond to the sampling location which the genomes were sourced from, not to the estimated species phylogeny of *Methanovorans*.

While all *Methanovorans* CIS loci share high amino acid identity, there are three subgroups of *Methanovorans* with distinct accessory genes and patterns of synteny in the CIS locus (Fig. 6). The amino acid identity of the CIS genes between *Methanovorans* within each subgroup is nearly 100%, while it is typically 75–85% between species in different subgroups. Interestingly, the *Methanovorans* species corresponding to each subgroup are not necessarily close phylogenetic relatives. Instead, CIS similarity appears to be driven by the geographic location at which the species were found, with one distinct subgroup at each of the Haakon Mosby Mud Volcano, Jaco Scar, and Scotian Basin sites (Figs. 6, S6). Interestingly, *Methanovorans* loci universally lack the ATPase gene generally associated with CIS, unlike the other genomes in this study, though knockout studies suggest this component is not required for CIS assembly and contraction (supp. text) (*35*).

### 3.4 Specialized nutrient acquisition pathways are sparse but gained by horizontal gene transfer in *Methanovorans*

Unlike many *Methanocomedenaceae* (ANME-2ab), *Methanogasteraceae* (ANME-2c), and methanogenic *Methanosarcinaceae*, most *Methanovorans* appear to lack the genetic potential to fix nitrogen. The exception to this pattern is the *Methanovorans* MAGs from the Scotian Basin site (sp. SB1, MAG S4 bin26 and MAG S7 bin36) (Figs. 7, S1). These two genomes each have a locus containing *nifHDK* and two small nitrogen regulatory genes. These genes are distinct from nitrogenase found in ANME-2 MAGs at the same site, though they have high sequence identity (*≈*90%). Gene phylogenies place these *Methanovorans* nitrogenases among the nitrogenases of *Methanocomedens* (ANME-2a) (Fig. S7). Protein alignments show a conserved deletion in ANME nifD, which corresponds to a missing loop on the protein surface above the FeMo cofactor active site in predicted protein structures (Fig. S8).

**Figure 7.**
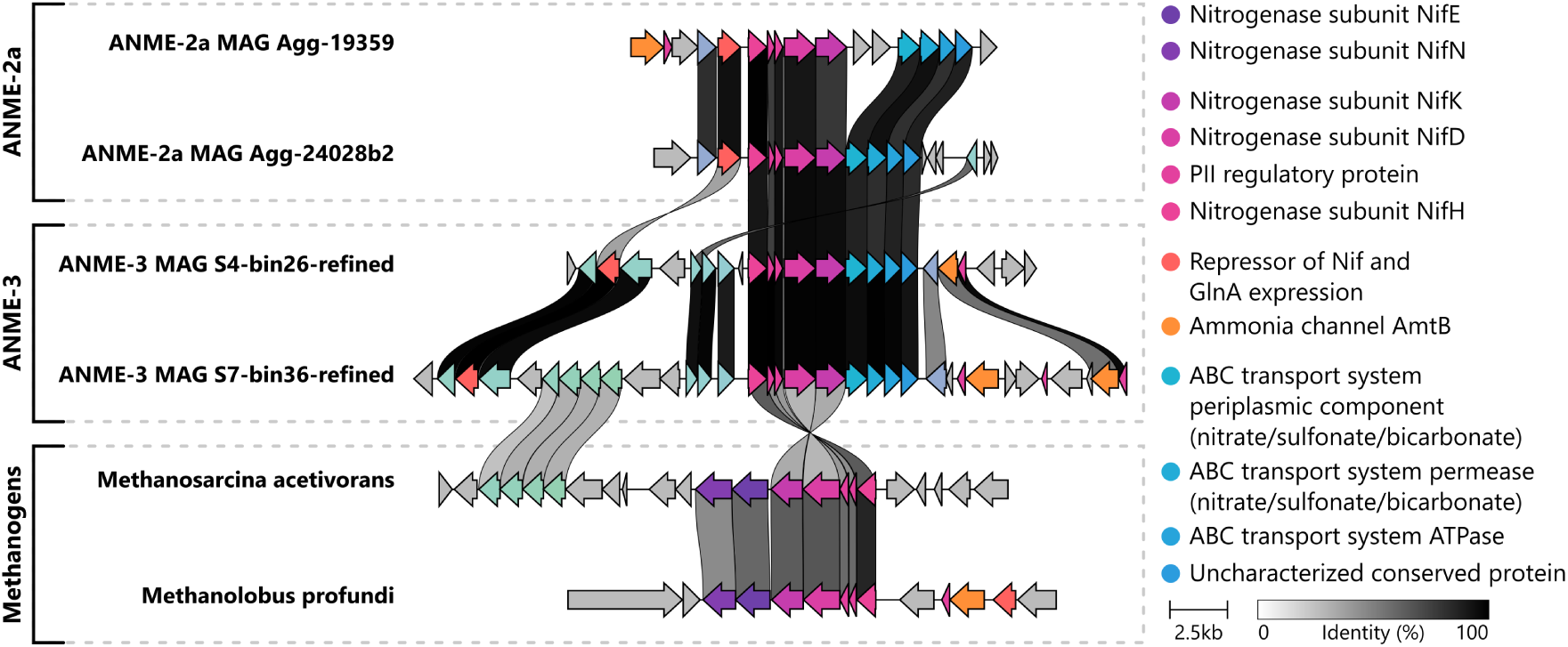
Representative nitrogenase loci of *Methanocomedens* (ANME-2a), *Methanovorans* (ANME-3), and methanogens. *Methanovorans* nitrogenase, which is present in only one *Methanovorans* species, shares very high sequence identity to nitrogenase from *Methanocomedens*. ANME nitrogenase loci have different gene content from methanogen nitrogenase loci. ANME appear to lack nifEN and contain additional putative ABC transporter components. Nif loci in all studied genomes were often flanked by conserved regulatory, transporter, or other genes, but *Methanovorans* nif loci appear to have reversed orientation relative to these flanking genes when compared to both *Methanocomedens* and methanogen loci.

The synteny of the *Methanovorans nif* locus also shares much higher similarity to the ANME-2 *nif* loci than to the loci of methanogens (Fig. 7). In place of *nifEN*, all ANME *nif* loci except ANME-2c MAG E20 have a conserved set of three ABC transporter genes (predicted to transport nitrate, sulfonate, or bicarbonate) and an uncharacterized conserved protein. This is unusual, as *nifEN* are considered essential for the formation of functional nitrogenase, though there may be exceptions (supp. text). There is also considerable variation in the genes surrounding the *nif* loci, but *Methanovorans* seem to have the highest similarity in these genes to *Methanosarcina*. However, the orientation of the locus in *Methanovorans* relative to the surrounding genes shared with both *Methanosarcina* spp. and ANME-2 is reversed (Fig. 7).

In addition to nitrogen fixation, some ANME may be able to use phosphonate instead of phosphate as a phosphorus source. Phosphonate catabolism is a moderately common microbial process in marine environments (*36*). Though substrate-specific pathways exist, the broad-spectrum metabolism is encoded as a pathway of 14 genes including C-P lyase which has been suggested to aid in adaptation to extreme phosphorus limitation (*36*). Nearly all *Methanovorans* genomes from Jaco Scar encode the complete C-P lyase system (Fig. S1). This is in stark contrast to the methanogen and most *Methanocomendaceae* and *Methanogasteraceae* genomes in this study, which never contain the full locus. Interestingly, one *Methanocomedens* (ANME-2a) and one *Methanogaster* (ANME-2c) genome in this study encoded a partial locus (Fig. S1).

## 4 Discussion

Since the initial 16S rRNA gene detection of *Methanovorans* (ANME-3) within methane seep environments, their high phylogenetic relatedness to methylotrophic methanogens has suggested that they diverged from their methanogen ancestors at a later point than any other known ANME clade based on 16S rRNA gene sequences (*37*, *25*). This close similarity has made *Methanovorans* an ideal candidate for deeper investigation of the evolutionary and genomic features responsible for the transition from methanogenic metabolism to syntrophic methanotrophy. Building from the two published genomes of *Methanovorans* (*16*), here we assembled genomic data of ten *Methanovorans* MAGs from three geographically distinct seep environments representing five species level clades.

Analysis of *Methanovorans* (ANME-3) genomes has shed new light on their evolutionary processes, but there is much about ANME left to uncover. In particular, we propose that *Methanovorans* may have begun adapting their central metabolic pathways to methanotrophy before acquiring complex EET machinery, which has intriguing implications for the evolution of ANME syntrophy with sulfate-reducing bacteria. As this EET machinery is thought to be important for ANME-SRB syntrophy, it is possible that the early ancestors of *Methanovorans* operated outside of any partnership coupled with reduction of metal oxides or humic acids, as has been suggested previously for other ANME clades (*38*, *39*) and observed in *Methanoperedens* (ANME-2d) (*40*, *41*). As some *Methanovorans* appear to be incompletely adapted to methanotrophy, this may also help explain the observation of many individual, partnerless *Methanovorans* cells at the Haakon Mosby Mud Volcano (*25*). However, it is worth noting that other ANME have also occasionally been observed outside of syntrophic partnerships as single cells (*4*, *42*, *43*) or monospecific aggregates (*4*, *44*, *45*, *46*), suggesting this may be a feature of all ANME in the right environmental conditions. Future studies of *Methanovorans* and the molecular basis for development and maintenance of ANME-SRB syntrophy will help confirm when in ANME evolution this partnership began.

### 4.1 The transition to methanotrophy in *Methanovorans* is ongoing

Though all *Methanovorans* originated from a common ancestor and are equally old, it appears that different lineages within *Methanovorans* are in different stages of losing pathways for methylotrophic methanogenesis (Fig. 2, 8, 9). Even in lineages which retain the genes for methylamine utilization and/or pyrrolysine biosynthesis, codon usage patterns suggest they are not likely to be expressed (Tab. S2). It is interesting that all *Methanovorans*, even those that retain pyrrolysineutilizing genes, have the same codon usage pattern as non-pyrrolysine utilizing organisms. Though selection is likely to be relaxed on the codon utilization and the biosynthetic genes on a similar timescale, changes in codon utilization require fewer mutations to become noticeable and thus this signal may precede any changes in biosynthetic genes when a non-canonical amino acid is lost. However, we still expect selection to be relaxed in these genes, which several recent studies suggest should lead to increased rates of gene loss (*47*, *48*, *49*). As expected from these results, we observe mutational disruptions and/or inactivating insertions in the *Methanovorans* MAG HMMV pyrrolysine biosynthesis locus. This suggests that we are observing *Methanovorans* while they are in the process of losing these genes and still optimizing for their new metabolism.

**Figure 8.**
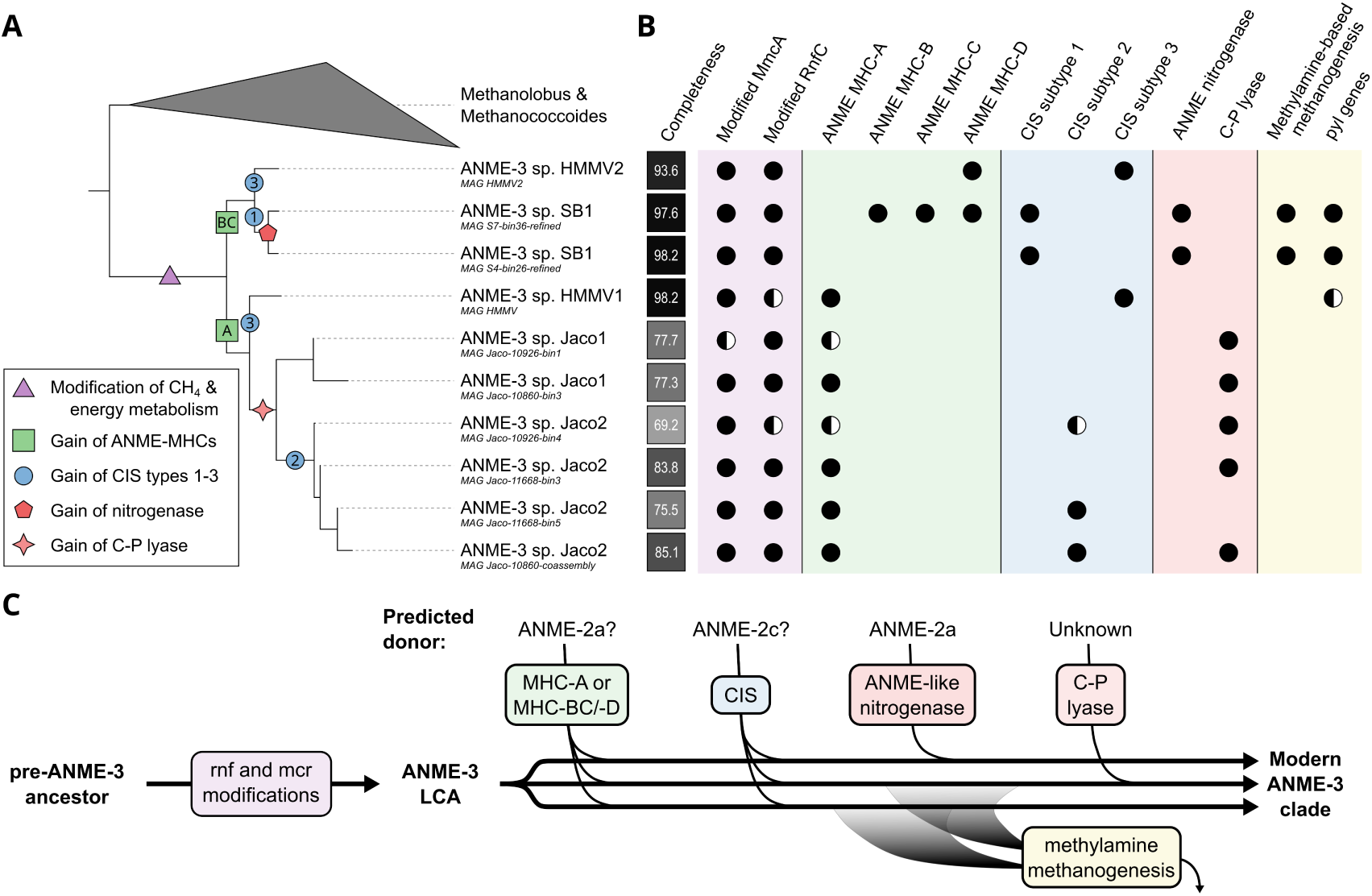
Proposed sequence of major events in *Methanovorans* (ANME-3) evolution. MmcA and RnfC are subunits of the energy conserving complex RNF. ANME MHCs are large multi-heme cytochromes proposed to facilitate direct intercellular electron transfer. CIS is a protein secretion system proposed to facilitate interspecies interactions between ANME-SRB syntrophic partners. Nitrogenase and C-P lyase are complexes which aid in nutrient acquisition. (A) Placement of evolutionary events on the estimated species phylogeny of *Methanovorans* based on the degree of conservation of genomic features in *Methanovorans*. Colored shapes indicate the timing of a given event, and text labels indicate distinct loci that may be involved in each event. (B) Presence and absence of distinctive features in *Methanovorans* genomes. Full circles indicate presence of the complete gene or locus while partially filled circles indicate presence of a recognizable fragment of the gene or locus. (C) Estimated timeline and provenance of distinctive features in *Methanovorans* genomes. Local modification of *mcr*, *rnf*, and *mmcA* genes and/or loci appear to have been present in the last common ancestor of the studied *Methanovorans* genomes. Acquisition of large ANME multi-heme cytochrome proteins likely occurred early after the diversification of the genus, followed by mainly vertical inheritance of these loci with some recombination. Acquisition of contractile injection systems, nitrogenase, and C-P lyase likely occurred late in the diversification of the genus, after global geographic dispersal.

**Figure 9.**
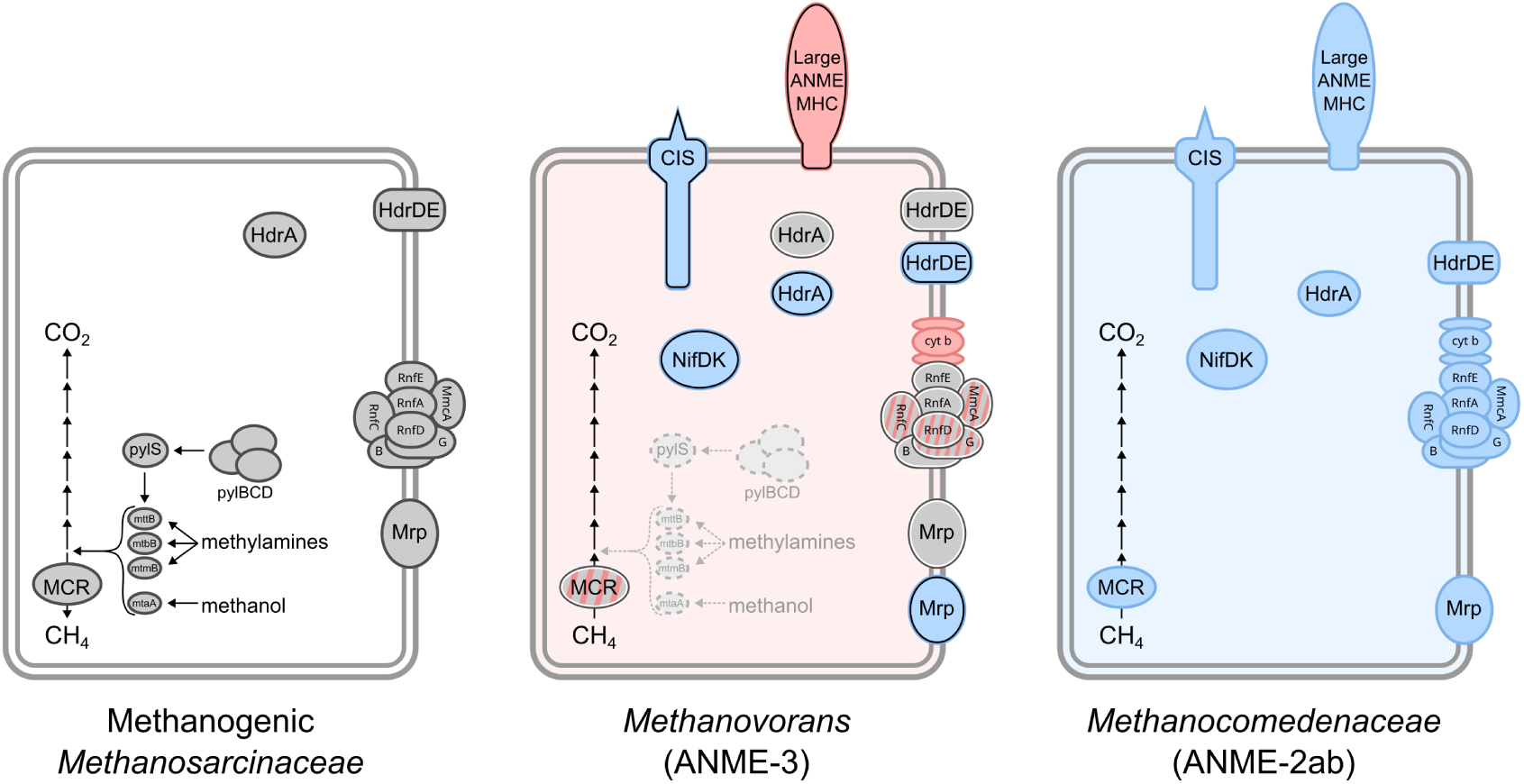
Comparison of gene content between representative examples of methanogenic *Methanosarcinaceae*, *Methanovorans* (ANME-3), and *Methanocomedenaceae* (ANME-2ab). Genes with a gray background are estimated to derive from methanogenic *Methanosarcinaceae* and genes in blue from *Methanocomedenaceae* in *Methanovorans*. Genes in red appear to be novel in *Methanovorans*. In *Methanovorans*, a black border indicates a gene is likely present due to horizontal gene transfer and a white border indicates presence via vertical inheritance. In central metabolic genes, *Methanovorans* appears to be a patchwork of vertically inherited methanogen derived genes and novel innovation. In accessory genes, *Methanovorans* appears to combine methanogen-like genes with genes derived from ANME-2 by horizontal transfer.

It appears that *Methanovorans* lineages are also in different stages of gene gain. The Mrp (Mnh) sodium-proton transporter and heterodisulfide reductase Hdr have several homologs, some of which are distinctive to ANME and others distinctive to methanogenic *Methanosarcinaceae* (File S2). We observed (supp. text) that some *Methanovorans* encode the methanogen-like homologs, some encode the ANME-like homologs, and several encode both (Fig. 9). This supports the hypothesis that the evolutionary transition of *Methanovorans* toward methanotrophy is ongoing. Also, though ANME-specific HdrA homologs have been proposed as a key metabolic innovation in ANME (*16*), the lack of conservation of these genes in *Methanovorans* suggests their acquisition is unlikely to be a precipitating event in the evolution of methanotrophy in *Methanovorans*. The similarity of *Methanovorans* with methanogens and their apparent recent divergence helps to explain why *Methanovorans* MAGs have not always been recognized as such. Most descriptions of *Methanovorans* (e.g. (*37*, *25*, *50*, *51*)) have relied 16S rRNA gene sequences to assign taxonomy, and thus MAGs which lack the 16S rRNA gene (like those from Dong *et al.* (*26*)) generally appear to be a methanogenic member of the *Methanosarcinaceae*. The introduction of the GTDB (*52*) has improved identification, with *Methanovorans* currently represented there by the g__DQIP01 annotation. The MAGs characterized in this study increase the known diversity within *Methanovorans* and further improve the identification of *Methanovorans* members based on additional marker genes. For example, we were able to use functional gene sequence similarity to identify a new candidate MAG from a sample from the Glendhu Ridge methane seep near New Zealand at 1974 m depth (NCBI: GCA 013139685.1) in the NCBI database that is likely to be a member of the *Methanovorans* genus.

### 4.2 Methanotrophy in *Methanovorans* was first shaped by convergent evolution

It has been clear since the first genomic investigations of ANME that they share the canonical 7-step methanogenesis pathway (*10*). This supported the lasting hypothesis that ANME accomplish the anaerobic oxidation of methane by ‘reverse methanogenesis,’ using the enzymes in the methanogenesis pathway in the opposite direction. However, aside from gene variations observed in *Methanophagales* (ANME-1) such as the lack of *mer* (*10*, *11*) and a distinct cofactor F430 and post-translational modifications in MCR (*53*), it has been difficult to discern key differences in this pathway between methanogens and ANME. Heterologous expression of MCR in methanogenic archaea has not clarified this situation. MCR subunits from ANME-1 do not complex with methanogenic subunits (*54*), and ANME-1 MCR may allow *Methanosarcina* to perform AOM (*20*). However, MCR from ANME-2 is highly similar to methanogenic MCR, to the point that subunits from ANME-2 are readily interchangeable with subunits from methanogenic MCR (*54*). The lack of a clear distinction between these enzymes has complicated attempts to understand the selective pressures acting on MCR in ANME as opposed to methanogens.

C1 metabolism in *Methanovorans* (ANME-3) appears to have undergone convergent evolution at the sequence level with *Methanocomedenaceae* (ANME-2ab) and *Methanogasteraceae* (ANME-2c), transitioning *Methanovorans* from a methanogen-like metabolism to an ANME-like metabolism. This is reflected in MCR gene phylogenies, which place *Methanovorans* between ANME-2 and methanogenic *Methanosarcinaceae*, unlike their position in universal marker gene phylogenies (Figs. 1, 3). This may be a phylogenetic signal suggesting there is a sequence-level distinction between ANME and methanogenic MCR independent of taxonomic lineage. Horizontal transfer of MCR has been previously shown across several archaeal lineages (*15*, *14*, *55*), and MCR can be hosted on borgs (*56*). However, the placement of *Methanovorans* as relatively equidistant from *Methanocomedenaceae*, *Methanogasteraceae*, and other *Methanosarcinaceae* is not consistent with horizontal transfer from other ANME. It is more plausible that MCR followed vertical inheritance in *Methanovorans*, as is typical for methanogenic and methanotrophic lineages (*57*), with positive selection (as observed in structural subunits McrABG) leading to the emergence of conserved features similar to *Methanocomedenaceae* and *Methanogasteraceae* at the sequence level. Our observation of positive selection in MCR genes during the evolution of *Methanovorans* is especially noteworthy considering previous observations that MCR is generally under tight purifying selection in methane seep environments (*58*). However, the residues which are conserved among ANME generally do not obviously interact with the active site or substrate binding channels in predicted structures of *Methanovorans* MCR, so their function is unclear (supp. text). Despite this, the universality of these conserved sequence features across *Methanovorans* suggests that MCR may have undergone convergent evolution prior to the last common ancestor of *Methanovorans* (Fig. 8).

Similar patterns of convergence are also observed in *Methanovorans* in the locus of the respiratory complex RNF, where the amino acid identity and the synteny conservation in adjacent genes between *Methanovorans* and other methanogenic *Methanosarcinaceae* is more consistent with convergence than with horizontal gene transfer. As with MCR, gene phylogenies and universal sequence features indicate that *Methanovorans* RNF loci have important commonalities with other ANME that are distinct from methanogenic *Methanosarcinaceae* and may have emerged before the last common ancestor of the genus (Fig. 8). This pattern of convergence is apparent in RNF subunits associated with electron transfer (RnfCDG) rather than ion translocation (RnfAE) (Fig. S3) (*59*). The RnfB subunit, which interacts with ferredoxin, does not show signs of convergence like other electron- handling subunits. However, this may simply indicate that the ferredoxin-RNF interaction is conserved across both ANME and methanogens, while the interactions of RNF with other electron carriers are not. Also, the convergent modification of heme-binding motifs in MmcA suggests ANME may require finely tuned redox environments in this protein and selection may be driving ANME MmcA towards similar electronic properties in each distinct lineage (supp. text). Overall, this suggests that the evolution of this locus may have facilitated innovations in electron flow through *Methanovorans* metabolism, possibly including enhancing their ability to perform extracellular electron transfer. This is further supported by recent studies which have suggested that while the multiheme cytochrome MmcA has a complex role in *Methanosarcina acetivorans*, it is important in energy conservation including facilitating intracellular and extracellular electron transfer in conjunction with RNF (*60*, *61*, *62*, *63*).

The convergent evolution of MCR and RNF in *Methanovorans* has interesting implications for the likely evolutionary timeline of ANME within the family *Methanosarcinaceae*. The early modification of these proteins suggests they may be of central importance in determining whether an organism is capable of gaining energy for growth through methanogenesis or methanotrophy. In addition, the convergent nature of these modifications suggests that ANME evolution may be triggered primarily by a change in environmental and/or ecological selective pressures. Such pressures could be introduced by the association with syntrophic partners (*9*), or by environmental conditions such as high methane concentrations in conjunction with metal oxide availability (*38*). It is conceivable and possibly expected that this new niche or selective pressure could result from a fortuitous horizontal transfer event of other genes like large multi-heme cytochromes, but we did not identify any other novel genome features thought to be important in ANME metabolism which appear to predate the last common ancestor of the *Methanovorans* genus. This leads us to conclude that these convergent modifications in energy and C1 metabolism were the initial step in the evolution of methanotrophy within the *Methanosarcinales*, and that the last common ancestor of the genus was therefore already metabolically committed to anaerobic methane oxidation. Though a recent study (*64*) has claimed to observe methanotrophic growth in methanogenic *Methanosarcinaceae*, we propose that these modifications may be important in tuning the metabolism of an organism for energy conservation and growth from anaerobic methane oxidation.

Convergent evolution has been observed in a wide diversity of microbial taxa during adaptation to a novel metabolism. Insect endosymbiont bacteria have converged on identical gene content, gene order, and/or metabolic capacity despite long divergence times or different taxonomic origin (*65*). Intracellular pathogens have convergently evolved mechanisms to survive within their host cells (*66*). Sulfide-oxidizing cable bacteria evolving from sulfate-reducing *Desulfobulbaceae* appear to have converged on similar electron flow pathways to some other sulfide-oxidizing bacteria and may employ electron conduit protein complexes which are non-homologous but structurally similar to those in *Geobacter* and *Shewanella* (*67*). Also, diverse sulfate-reducing bacteria have converged on similar extracellular electron transfer pathways when adapting to syntrophy with ANME (*9*). However, *Methanovorans* is notable in that their convergence with other ANME is not only at the level of gene content or genome organization, but also in highly similar modifications at the sequence level to critical metabolic genes. It has been suggested that parallel evolution at the amino acid level may occur due to a drive to maximize adaptation while minimizing pleiotropic effects (*68*). Striking examples in *Methanovorans* are the parallel evolution of a CIDCH heme-binding motif in the MmcA protein which is conserved among ANME (Fig. S5) or the high conservation of certain residues in ANME MCR not shared with methanogens. Given the highly conserved nature of MCR and heme-binding environments, this suggests these protein regions may be under precise selective pressures in ANME which are distinct from the needs of methanogens. These protein regions also present potential targets for genetic modification in closely related methanogens (*69*) to more definitively test these hypotheses and mechanistic details of AOM.

### 4.3 Horizontal gene transfer drives later optimization and specialization in Methanovorans

Many distinctive ANME-like features vary in their gene sequences and presence or absence in *Methanovorans*, including energy metabolism genes like the large ANME multi-heme cytochromes, ANME-like Hdr, and ANME-like Mrp/Mnh, export systems like contractile injection systems (CIS) and a putative lipoprotein export system, and nutritional genes like nitrogenase and C-P lyase (supp. text, Fig. 9). Based on sequence similarity of ANME genes to other taxa, the polyphyly of gene sequences, synteny of surrounding genes, and patterns of presence and absence in *Methanovorans* genomes, each of these features appear to be the result of horizontal gene transfer in *Methanovorans*. While it is possible the genus had broad acquisition of many or all of these systems followed by differential loss, this is not a parsimonious explanation of the observed patterns.

The pattern of gene presence and absence also suggests that the MHCs were relatively early acquisitions within *Methanovorans*, while CIS, nitrogenase, and C-P lyase were acquired later, likely in response to local environmental conditions or specific syntrophic partner associations (Fig. 8). In the case of the MHCs, there are two deeply branching *Methanovorans* subclades with different complements of MHC loci, even when members of each subclade were found in the same sampling location. This points to the conclusion that none of these MHC genes were present in the common ancestor of *Methanovorans* but were acquired relatively early in the diversification of the genus, potentially before their widespread geographic dispersal. Also, the existence of a fusion between MHC-B and MHC-C in *Methanovorans* suggests that these proteins may function together in other ANME (supp. text).

Unlike the MHCs, loci such as nitrogenase and C-P lyase are found only in small groups of closely related MAGs from single sampling sites. Also, the nitrogenase genes found in *Methanovorans*, like those in other ANME (*70*), belong to a separate clade than genes from related methanogens and other archaea (Fig. S7). Though the CIS is more widespread within *Methanovorans*, the distribution of CIS subtypes also suggests that these systems were acquired independently at each of the sampled geographic locations after global dispersal of the genus. Overall, limitation of these genes to specific locales suggests that these late-acquisition loci may play a role primarily in optimization to specific environmental or ecological conditions, and supports the hypothesis of Ruff *et al.* (*5*) that even when observing globally distributed taxa, methane seep microbiota are highly shaped by local diversification.

The contractile injection system (CIS) in *Methanovorans* is a particularly instructive example of local specialization in ANME. Their phage-like protein translocation system (PLTS) is a class of CIS which is structurally similar to other systems like the type VI secretion system but share only distant evolutionary roots (*71*, *72*). Many functions of CIS have been proposed, including toxin delivery (*73*, *74*, *75*), intercellular communication and quorum sensing (*73*, *76*), biofilm restructuring (*76*), facilitation of horizontal gene transfer (*75*, *77*), and outer membrane vesicle recruitment (*77*). While the function of CIS in ANME is not well understood, prior studies have proposed a role in influencing the specificity of the ANME-SRB partnership (*16*). A similar role has been proposed for the analogous type VI secretion system found in many SRB (*9*). The distinct CIS subtypes observed in *Methanovorans* from each geographic site may potentially represent adaptations to differences in the syntrophic partnerships available at those sites. For example, while *Methanovorans* were observed in partnership mainly with *Desulfobulbaceae* at the Haakon Mosby Mud Volcano (*25*), it has been proposed that they may partner with members of the SEEP-SRB1a clade at other sites (*78*, *8*) (supp. text). This geographic variation in CIS subtypes among *Methanovorans* is also notable because the Haakon Mosby Mud volcano has only been active for about 30 ky (*79*), which could indicate that the acquisition of a location-specific CIS subtype and possibly other features can happen over relatively short geologic timescales in ANME evolution.

The acquisition of CIS may also be an example of convergence in multispecies interactions, despite being different from the convergent parallel evolution discussed in the context of MCR and RNF. It has been proposed that multispecies interactions may exhibit convergence in a variety of factors including morphology and functional interactions (*80*). Though the function of CIS in ANME is not known, a recent paper demonstrated that CIS are upregulated in *Geobacter metallireducens* when performing direct extracellular electron transfer (DIET) involving physical association with *Geobacter sulfurreducens* and *Methanosarcina barkeri*, resulting in an increased lag time to establish DIET (*81*). This effect was reduced by the addition of conductive carriers like the quinone AQDS. Previous transcriptomic work suggests that ANME may also differentially express CIS components in response to the addition of AQDS to decouple them from their syntrophic partners (*82*). Though CIS are not present in all organisms that participate in DIET, the potential role of CIS in modulating some DIET syntrophies is worthy of future study.

The ANME MHC and CIS gene loci are quite large (over 25 kb and 17 kb respectively in *Methanovorans*), raising the question of the mechanics of their transfer. While these alone are larger than many typical plasmids and viral genomes, megaplasmids have been reported in *Methanosarcinales* (the taxonomic order containing the organisms in this study) and are common in *Haloarchaea* (*83*). Additionally, extrachromosomal elements up to 1 Mb known as borgs (*56*) and unusually large plasmids up to 200 kb (*84*) have recently been identified in *Methanoperedens* spp. (ANME-2d) and proposed to act as gene transfer agents, including of large MHCs. However, such extrachromosomal elements have yet to be identified in the ANME clades studied here. Further investigation into mobile genetic elements in *Methanovorans* is likely needed to fully understand the acquisition of these large gene cassettes, but there is precedent for large potential gene transfer agents in related archaea.

### 4.4 ANME evolution may involve adaptation to nutrient limitation

Several of the common ANME adaptations may be responses to increased nutrient limitation, as their methane seep habitat is often characterized by steep geochemical gradients, and aspects of ANME-SRB syntrophy are known to vary with nutrient concentrations (*85*). Perhaps the most notable such adaptation is the acquisition of nitrogen fixation genes. Though ANME nitrogenase genes are distinct from most characterized sequences, they are known to be functional as ANME fix nitrogen under natural conditions (*70*, *86*). The nitrogenase sequences found in this study are highly similar to characterized ANME nitrogenase genes. Based on gene phylogeny and locus synteny, it seems most likely that the *Methanovorans* from the Scotian Basin have acquired their nitrogenase horizontally from *Methanocomedens* (ANME-2a), but may have integrated it at a similar genome position to the locus in their *Methanosarcinaceae* relatives. It is unclear why the Scotian Basin *Methanovorans* lineages and many ANME-2 have acquired nitrogenase, but those from HMMV and Jaco Scar lack it. Ammonia is present at relatively high concentrations in the subsurface-derived fluids at the HMMV (*87*) and not measured at the carbonate-hosted Jaco Scar site, so this lack of nitrogenase may be due to specific conditions of the sampling sites.

Another potentially site-specific nutritional adaptation is the acquisition of phosphonate metabolic genes by several species of *Methanovorans* localized to the Jaco Scar site. The ability to use phosphonate has not been described in any other methanotrophic lineage to date, though C-P lyase has been found in other archaeal species (*88*). C-P lyase has broad substrate specificity but is a relatively expensive complex to maintain, requiring expression of a locus of 14 genes (*36*). For this reason, this broad-specificity phosphonate metabolism is most abundant in regions of the ocean with severe phosphorus limitation (*36*). While most surveys of the correlation between C-P lyase and nutrient conditions have focused on the pelagic environment, it is possible that a similar dynamic could be at play in the sediments. Interestingly, we also observed part of the C-P lyase locus in a *Methanocomedens* (ANME-2a) and a *Methanogaster* (ANME-2c) genome, (Fig. S1), and BLAST searches indicated that several other publicly available ANME genomes also encode C-P lyase with typical amino acid identity of at least 80% to the *Methanovorans* sequences. This suggests that this metabolism could be more widespread across ANME clades than is currently recognized.

The enrichment of toxin-antitoxin systems and transporter complexes in ANME genomes relative to other *Methanosarcinaceae* also may be a response to nutrient limitation, though their role in ANME is not currently well characterized. For example, the toxin-antitoxin (TA) systems *relBE* and *mazEF*, which are enriched in ANME genomes, have been implicated in the response to nutritional stress in bacteria (*89*). In that context, these TA systems may have a reversible bacteriostatic effect, allowing the cell to go dormant until nutrient availability increases. The observed enrichment in metal ion transporters may relate to the high trace metal requirement expected for for methanotrophy (*90*, *91*). However, it is not clear why ANME would have higher trace metal requirements than related methanogens (*90*). ANME enrichment in branched-chain amino acid transporters may be related to the growth advantage in stationary phase (GASP) phenotype, helping take advantage of necromass in the environment (*92*, *93*). ANME are estimated to incorporate only 1% of the methane they oxidize into biomass (*94*), so while it is unlikely that ANME would metabolize necromass heterotrophically, the ability to supplement autotrophic biosynthesis with existing amino acids would likely be advantageous. Similar potential for organic molecule assimilation has also been proposed in *Ca.* Methanophagales (ANME-1) on the basis of isotope labeling (*95*). Construction of extracellular barriers, as seen in ANME aggregates, can also be a response to stationary phase in bacteria (*96*) or to various stresses in archaea (*97*). While not conclusive, these observations suggest that the role of nutrient limitation in shaping ANME ecology is worthy of future study.

### 4.5 Outlook

Our proposal that the early ancestors of *Methanovorans* primarily underwent gradual convergent evolution also has implications for the ecology of methanotrophy in natural environments. It suggests that some methanogenic archaea such as the early ancestor of *Methanovorans* may oxidize methane under specific environmental conditions. In fact, one study has claimed to observe growth under AOM in *Methanosarcina* (*64*), but our results suggest that the reversibility of enzymes in the methanogenesis pathway is not sufficient to evolve methanotrophic metabolism, and there must also be energetic innovations to couple AOM to growth. Overall, this suggests that it should be possible for current methanogenic lineages to give rise to novel ANME lineages in the right environment, though what conditions might be necessary for this remains unclear. It is worth noting that the methylotrophic methanogenesis pathways common to the *Methanosarcinaceae* family already run several steps from the canonical hydrogenotrophic methanogenesis pathway in reverse (Fig. 9), so these organisms may require fewer adaptations to fully reverse the pathway as seen in ANME. Prior studies have reported the capacity for methane oxidation in methylotrophic *Methanosarcina* species under laboratory conditions (*20*, *21*, *22*, *23*), and genetic manipulation in related species (*69*) of the genes identified here may help test our hypotheses. However, understanding the potential for methanogens to perform AOM in natural environments will require additional ecological study.

The relative recency of the emergence of *Methanovorans* may also facilitate study of the biogeography of methane seep ecosystems. Ruff *et al.* (*5*) proposed that methane seep microbiomes are shaped by global dispersal and local diversification based on site-specific environmental conditions. While our data broadly suggest that *Methanovorans* may exemplify both of these processes, we do not have sufficient data to confidently assess their biogeography. However, our observation of their potentially rapid and recent diversification at new seep sites like the Haakon Mosby Mud Volcano suggests that *Methanovorans* could be a useful case study through which to examine the processes shaping dispersal and diversity of methane seep taxa. Future analysis of a greater diversity of *Methanovorans* MAGs from new sampling sites such as the Glendhu Ridge MAG mentioned above and recently published MAGs from a terrestrial mud volcano (*50*, *51*) may clarify how such taxa spread and adapt to each new ecosystem they encounter.

## 5 Methods

### 5.1 Sample collection

Authigenic carbonate samples were collected from the Jaco Scar hydrothermal seep located on the Costa Rica margin at approximately 1800 m depth using the crewed submersible *Alvin* during research expedition AT42-03 on *R/V Atlantis*. All rock samples were collected at a site with an overlying water temperature of 6 *^◦^*C and near shimmering water, indicating local discharge of hydrothermal fluid. Rock sample 10860 (source of *Methanovorans* (ANME-3) MAGs Jaco-10860-bin3 and Jaco-10860-coassembly) was collected on dive AD4972 on 18 October, 2018 at coordinates 09*^◦^* 07.067*^′^*N, 84*^◦^* 50.3707*^′^* W at a water depth of 1784 m. This rock was associated with a pink microbial mat which was preserved separately. Rock sample 10926 (source of *Methanovorans* (ANME-3) MAGs Jaco-10926-bin1 and Jaco-10926-bin4) was collected on dive AD4973 on 19 October, 2018 at coordinates 09*^◦^* 07.070*^′^* N, 84*^◦^* 50.3706*^′^*W at a water depth of 1784 m. Rock sample 11668 (source of *Methanovorans* (ANME-3) MAGs Jaco-11668-bin3 and Jaco-11668-bin5) was collected on dive AD4989 on 4 November, 2018 at coordinates 09*^◦^* 07.067*^′^* N, 84*^◦^* 50.367*^′^* W at a water depth of 1738 m. All samples were processed and frozen on board at *−*80 *^◦^*C until later DNA extraction in the laboratory.

### 5.2 DNA extraction and sequencing

A 250 mg subsample of carbonate rock from each of the samples 10860, 10926, 11668, and the mat associated with rock 10860 was crushed into a powder with a sterile mortar and pestle and DNA was extracted from each of the mineral samples following previously published protocols (*98*). Metagenomic sequencing of the genomic DNA extracts was outsourced to Quick Biology (Pasadena, CA, USA) as previously described (*99*). Complete details for the methods are also provided in the supplement.

### 5.3 Metagenomic analysis and assembly of new ANME genomes

The sequencing reads were assembled using SPAdes v3.12.0 (*100*, *101*). For samples 11926 and 11668 libraries were assembled individually, while those from samples 10860 and 10860 mat were assembled in a coassembly. From the de-novo assemblies, we performed manual binning using Anvio v6 (*102*). We assessed the quality and taxonomy affiliation from the obtained bins using GTDB-tk v1.5.0 (*52*, *103*) and checkM v1.1.3 (*104*). Genomes affiliated with *Methanovorans* (ANME-3) were further refined via a targeted-reassembly pipeline as previously described (*99*).

### 5.4 Reference genome selection, refinement, and taxonomy

We downloaded all 449 genomes and MAGs (metagenome assembled genomes) listed as part of the class *Methanomicrobia* in the NCBI database as of June 3, 2019 (*105*). We used our inhouse database to supplement with 58 previously published MAGS from *Methanocomedens Methanomarinus*, and *Methanogaster* (ANME-2a, -2b, and -2c, respectively) (e.g. (*16*)). The inhouse database also included (at the time) unpublished MAGs, including *Methanocomedens* (ANME-2a) single aggregate genomes later published by Yu *et al.* (*106*) and two *Methanovorans* (ANME-3) MAGs derived from unassembled sequence data from Ruff *et al.* (*87*) and later published by Chadwick *et al.* (*16*). In addition, two MAGs from the Scotian Basin published by Dong *et al.* (*26*) were identified for further analysis. Dong *et al.* (*26*) reported a relatively high abundance of *Methanovorans* (ANME-3) at this site, and assembled two high-quality MAGs which clustered within the family *Methanosarcinaceae*, labeled S4 bin26 and S7 bin36. From their position in the CheckM reference tree, it appeared that these two Scotian Basin MAGs were unclassified members of *Methanovorans* (ANME-3). We manually refined these MAGs using the Anvi’o platform by examining coverage and gene taxonomy across their contigs (*107*).

These genomes and MAGs were analyzed with CheckM version 1.1.2 using the lineage workflow (*104*). Only high quality genomes with *≤* 500 scaffolds, an N50 *≥* 10 kb, and an estimated quality (completeness *−* 5 *×* redundancy, defined by Parks *et al.* (*108*)) *≥* 50 were retained for analysis. A further eleven genomes were excluded because they clustered outside the *Methanomicrobia* in the CheckM reference tree, and four genomes were excluded because they seemed to have many frameshifted proteins. This left 365 of the NCBI genomes and MAGs and 34 of the internal ANME MAGs. In addition to these genomes and MAGs, four of the new Jaco Scar *Methanovorans* assemblies also met all of the quality criteria. The remaining two (MAG Jaco-10926-bin1 and MAG Jaco-10926-bin4) met all criteria except they had a scaffold N50 less than 10 kb but were nevertheless retained for further analysis (Tab. S1, file **??**).

We selected a subset of genomes providing a representative sample of each major clade across the *Methanocomedens* (ANME-2a), *Methanomarinus* (ANME-2b), *Methanogaster* (ANME-2c), and the *Methanosarcinaceae* (Tab. S1). When possible, each genus was represented by multiple genomes. Preference was given to NCBI type genomes and genomes with high estimated quality scores. Together with the six new assemblies from Jaco Scar, this resulted in a final set of 39 genomes.

The 39 study genomes were analyzed with GTDB-Tk version 1.5.0 using GTDB release R06-R202 and the classify wf command (*103*, *52*). Genomes with *≥* 95% ANI, as computed by PyANI (*109*) ANIb (*110*), were considered to represent the same species.

### 5.5 Gene calling and annotation

The 39 study genomes were analyzed using the Anvi’o platform version 7.1 (*102*). Gene calling was performed by Prodigal v2.6.3 (*111*). Canonical tRNAs were identified using tRNAscan-SE v2.0.7 (*112*). Sequences for the tRNA *pylT* were manually identified in some study genomes using published *pylT* sequences and genome coordinates (*28*, *113*). Additional *pylT* sequences were found iteratively with blastn v2.10+ (*114*) using the published sequences to query against the study genomes, then using the resulting sequences as a new query, and so on. This approach failed to identify the *pylT* gene in some of the methanogen genomes with otherwise complete *pyl* pathways, so this was followed by applying a custom HMM based on the collected sequences built with HMMER v3.3.1 (*115*).

Genes were annotated within Anvi’o using the COG database (*116*), Pfam version 32.0 (*117*), and KO-fam (*118*). Gene annotations were separately performed with eggNOG-mapper version 2.0.1 (*119*, *120*) and imported into Anvi’o. For analyses of RnfC and large multi-heme cytochrome proteins, protein domains were identified with InterProScan 5 (*121*, *122*). Putative *c*-type cytochrome proteins were identified by searching for the typical CXXCH heme binding motif in the protein sequences of each organism using the scripts prot motif search.pl and prot motif count.pl at https://github.com/orphanlab/anme3evo-code. Codon frequency analyses were performed using the Anvi’o command anvi-get-codon-frequencies.

Conserved ANME-specific amino acid residues in methyl-coenzyme M reductase subunits and phylogenetic marker gene sequences were identified using the script filter-alignment.py which can be found at https://github.com/orphanlab/anme3evo-code. Briefly, this script identifies positions in an amino acid alignment where the “ingroup” sequences meet the specified conservation thresholds and the ingroup consensus sequence is never observed in the outgroup at that position. The “ingroup” parameter included all ANME gene sequences and the “outgroup” parameter included all non-ANME sequences. The minimum identity threshold was set to 0.6 or 0.7, the maximum allowed gaps was 0.3, and the property conservation threshold was set to 0. Because the number of conserved residues was very small, confidence intervals were calculated for the number of conserved residues in each gene assuming a Poisson distribution. These bounds were then normalized by dividing by the sequence length to obtain bounds on the rate of conserved residues per thousand amino acids.

### 5.6 Phylogenetic analysis

All gene alignments were performed with MUSCLE v3.8.1551 (*123*). Each of the alignments was trimmed with the trimAl software version 1.4.rev15 using the -automated1 setting (*124*). Construction of the trees was performed with RAxML v8.2.12 (*125*). The specific settings used for each tree are listed in file S3. Phylogenetic trees were visualized using iTOL (*126*).

Genes from the Anvi’o marker set Archaea 76 (*127*) and the 16S rRNA genes were extracted from the genomes using HMMER (*115*). Only those marker genes which were present in at least 90% of the genomes (36 of 39) were used for further analysis. These marker genes were aligned using MUSCLE (v3.8.1551) and the alignments were concatenated using Anvi’o.

The methyl-coenzyme M reductase (*mcr*) and Rhodobacter nitrogen fixation (*rnf*) gene sequences were extracted from the genomes on the basis of functional annotation. One sequence annotated as methylcoenzyme M reductase subunit A (*mcrA*) was discarded because it was highly divergent from the other sequences. The genes of the contractile injection system (CIS) were extracted from the genomes on the basis of functional annotation, informed by synteny (i.e. gene order and orientation in a locus) and sequence identity to known CIS genes (*35*, *71*). Each gene was aligned individually and a concatenation was made with the genes present in at least 13 of the 15 identified CIS loci.

Testing for positive selection was performed using the aBSREL method of HyPhy v2.5.50 (*128*). The inputs were the methyl-coenzyme M reductase subunit A (McrA) codon alignment and the concatenated marker gene phylogeny (pruned to only include taxa encoding a complete *mcrA* gene). The codon alignment was generated with PAL2NAL v14 (*129*). The only branch tested for positive selection was the branch leading to the last common ancestor of *Methanovorans* (ANME-3).

Trees were also constructed using marker gene sets from Phylosift (*130*) and single-copy core genes from the pangenome analysis, but these did not provide high support for all branches.

### 5.7 Pangenome and functional analysis

The 39 genomes were each placed into one of three groups: ANME-2 (11 genomes), ANME-3 (10 genomes), or Methanogens (17 genomes). These groups were used for several downstream analyses.

The completeness of metabolic pathways in the 39 genomes was assessed with KEGG Decoder version 1.2 (*131*). Enrichment of various metabolic pathways among the three groups was assessed with the Anvi’o metabolic enrichment pipeline (*132*).

We constructed a pangenome database using the Anvi’o pangenome workflow (*133*). Briefly, this involved running an all-vs-all DIAMOND-sensitive v2.0.6 (*134*) search of the protein sequences in the study genomes, then performing a clustering analysis using the MCL algorithm (*135*) using the default settings. This database was then used to analyze the enrichment of functional annotations among the three groups using the Anvi’o functional enrichment pipeline (*132*). Enrichment was computed for COG functions, KOfams, Pfams, EggNOGs, and pangenome protein families (IDENTITY in Anvi’o). Examining the enrichment of the pangenome protein families enabled us to distinguish between multiple proteins with the same annotation in the case that only a subset of those was ANME-specific, or to resolve an ANME-specific protein that was annotated differently by each database.

The standard Anvi’o enrichment pipelines assess enrichment by comparing the proportion of genomes in each group which have a certain annotation present. If the annotation is present multiple times per genome, this information is not used. We modified this pipeline to compute enrichment by comparing the frequency of an annotation within the genomes in each group, and used this to assess the enrichment of COG categories among the three groups (inspired by Allen *et al.* (*136*)). Frequencies were computed based on the total number of annotations per group (e.g. a frequency of 0.082 for COG category C — energy production and conversion — in the ANME-2 group indicates that 8.2% of the combined COG annotations from all 11 ANME-2 genomes fell into category C). This script is available as anvi-script-frequency-enrichment.sh at https://github.com/orphanlab/anme3evo-code.

The prevalence of multi-heme cytochromes (MHCs) in the study genomes was compared using one-way ANOVA, with the genomes grouped as ANME-2, ANME-3, or Methanogens, as above. Post-hoc pairwise comparisons between the three groups used 2-tailed Student’s t-tests with Bonferroni’s correction. Each analysis used 0.05 as the significance level. This process was used to compare the total number of *c*-type cytochromes in the genomes, the number of single-heme proteins, and the number of multi-heme proteins (proteins with two or more heme binding motifs).

### 5.8 Synteny analysis

We looked for potential instances of HGT between ANME groups by finding regions of identical synteny (identical gene order and orientation between loci) between *Methanovorans* (ANME-3) and other ANME. Briefly, gene neighborhoods between 4 and 15 genes long were extracted from the study genomes using the Anvi’o command anvi-analyze-synteny. The genes in the extracted neighborhoods were named using the protein family IDs from the pangenome analysis. We then used a custom script to group together identical gene neighborhoods across the study genomes, and to attach functional annotation information to these neighborhoods. This script is available as anvi-script-extract-synteny-groups.sh at https://github.com/orphanlab/anme3evo-code. Visualization and comparison of gene neighborhoods was performed using Artemis version 18 (*137*) and clinker version 0.0.21 (*138*).

### 5.9 Protein structure prediction

Structural prediction of proteins was done using AlphaFold 2 (*139*). Charge surface calculations were performed with APBS v3.0.0 and PDB2PQR v3.5.1 using the public web server (*140*). Visualization of the predicted structures was done using UCSF ChimeraX v1.6.1 (*141*, *142*).

## 6 Acknowledgments

We would like to thank Ranjani Murali and Grayson Chadwick for informative discussions regarding this work. This report was prepared as an account of work sponsored by an agency of the United States Government. Neither the United States Government nor any agency thereof, nor any of their employees, makes any warranty, express or implied, or assumes any legal liability or responsibility for the accuracy, completeness, or usefulness of any information, apparatus, product, or process disclosed, or represents that its use would not infringe privately owned rights. Reference herein to any specific commercial product, process, or service by trade name, trademark, manufacturer, or otherwise does not necessarily constitute or imply its endorsement, recommendation, or favoring by the United States Government or any agency thereof. The views and opinions of authors expressed herein do not necessarily state or reflect those of the United States Government or any agency thereof.

## 6.2 Author contributions

- Conceptualization: PHW, DRS, VJO
- Data curation: PHW, RLP, DRU
- Funding acquisition: VJO
- Investigation: PHW
- Methodology: PHW, DRS
- Project administration: VJO
- Resources: VJO
- Software: PHW
- Supervision: VJO, DRS
- Visualization: PHW, SER
- Writing – original draft: PHW
- Writing – review & editing: PHW, DRS, RLP, DRU, SER, VJO

## 6.4 Data and materials availability

All genome data used in this study can be found at DOI 10.22002/hgawh-n7907 along with paired annotation files. The newly assembled genomes are deposited with NCBI under (BIOPROJECT ID). All original code by the researchers used in this work is available at https://github.com/orphanlab/anme3evo-code. All other data are available in the main text or the supplementary materials.

## 7 Supplementary Material

### 7.1 Methods

#### 7.1.1 DNA extraction and sequencing

A 250 mg sample of rocks 10860, 10926, 11668, and the mat associated with rock 10860 was crushed and DNA was extracted from each of the mineral samples following previously published protocols (*98*). Metagenomic analysis from the extracted genomic DNA was outsourced to Quick Biology (Pasadena, CA, USA) for library preparation and sequencing as previously described (*99*).

Libraries were prepared with the KAPA Hyper plus kit using 10 ng of DNA as input. This input was subjected to enzymatic fragmentation at 370C for 10 min. After end repair and A-tailing, the DNA was ligated with an IDT adapter (Integrated DNA Technologies Inc., Coralville, Iowa, USA). Ligated DNA was amplified with KAPA HiFi HotStart ReadyMix (2x) for 11 cycles. Post-amplification cleanup was performed with 1x KAPA pure beads. The final library quality and quantity were analyzed and measured by Agilent Bioanalyzer 2100 (Agilent Technologies, Santa Clara, CA, USA) and Life Technologies Qubit 3.0 Fluorometer (Life Technologies, Carlsbad, CA, USA) respectively. Finally, the libraries were sequenced using 150 bp paired-end reads on Illumina HiSeq4000 Sequencer (Illumina Inc., San Diego, CA). After sequencing, primers and adapters were removed from all libraries using bbduk78 (https://sourceforge.net/projects/bbmap/) with mink=6 and hdist=1 as trimming parameters, and establishing a minimum quality value of 20 and a minimal length of 50 bp.

#### 7.1.2 Metagenomic analysis and genome assembly

The sequencing reads were assembled using SPAdes v. 3.12.0 (*100*, *101*). For samples 11926 and 11668 libraries were assembled individually, while those from samples 10860 and 10860 mat were assembled in a coassembly. From the de-novo assemblies, we performed manual binning using Anvio v. 6 (*102*). We assessed the quality and taxonomy affiliation from the obtained bins using GTDB-tk (*103*, *52*) and checkM (*104*). Genomes affiliated to *Methanovorans* (ANME-3) were further refined via a targeted-reassembly pipeline as previously described (*99*).

In this pipeline, the original reads were mapped to the bin of interest using bbmap (https://sourceforge.net/projects/bbmap/), then the mapped reads were assembled using SPAdes and finally the resulting assembly was filtered discarding contigs below 1500 bp. This procedure was repeated during several rounds (between 16-18) for each bin, until we could not see an improvement in the bin quality. Bin quality was assessed using the checkM and considering the completeness, contamination (*<*5%), N50 value and number of scaffolds. The resulting bins were considered as metagenome-assembled genomes (MAGs).

### 7.2 Supplemental Results and Discussion

#### 7.2.1 Partnership between *Methanovorans* and sulfate-reducing bacteria

At the Haakon Mosby Mud Volcano, *Methanovorans* (ANME-3) were described in aggregate partnerships with relatives of the genus *Desulfobulbus* and sometimes with an unidentified bacterial partner, unlike other ANME (*25*). Based on 16S taxonomy data from the Jaco Scar and Scotian Basin (*26*) sites, *Desulfobulbus* does not seem to have a significant presence at the depth horizons where *Methanovorans* were present. However, members of the SEEP-SRB1 clade, of which SEEP-SRB1a has been proposed as a potential partner for *Methanovorans* (*8*, *78*), were present in moderate abundance at the depth horizons containing *Methanovorans*. If SEEP-SRB1a are the primary partner for *Methanovorans* this could be a potential driver for convergent evolution with other ANME, as this clade is also the primary partner for *Methanocomedens* (ANME-2a) and *Methanogaster* (ANME-2c) (*78*). It is also interesting to consider whether incomplete adaptation to such a syntrophy may help explain why *Methanovorans* have also been observed in partnership with a variety of organisms as mentioned above, or as single cells with no apparent SRB partner (*25*).

#### 7.2.2 ANME are enriched in mobile genetic elements, defense mechanisms, and multi-heme cytochromes

Analysis of gene and functional enrichment profiles between *Methanocomendaceae* (ANME-2ab) and *Methanogasteraceae* (ANME-2c), *Methanovorans* (ANME-3), and methanogenic *Methanosarcinaceae* revealed several gene families or functions which seem characteristic of ANME. For some of these, such as members of the peptidase U32 or Roadblock GTPase modulator superfamily (file S1), it is difficult to interpret how they might relate to other aspects of ANME metabolism. However, some patterns do emerge when examining enrichment of mobile elements, defense mechanisms including toxin-antitoxin systems, and multi-heme cytochrome proteins.

Analysis of the frequencies of each COG category in *Methanocomedenaceae* and *Methanogasteraceae* and *Methanovorans* indicated that ANME are significantly enriched in mobile elements (X) and defense mechanisms (V) relative to methanogenic *Methanosarcinaceae* (file S1). Relative to other *Methanosarcinaceae*, ANME are enriched in genes annotated with COG categories V (defense mechanisms) and X (mobile elements), but relative to the outgroup *Methanothrix*, ANME are only enriched in the latter. Despite a lack of enrichment in COG category V relative to *Methanothrix*, the elevated abundance of these genes in *Methanovorans* over other *Methanosarcinaceae* suggests the genes could still be relevant in ANME evolution. A large component of the defense mechanism category in ANME appears to be an abundance of toxin-antitoxin (TA) systems, which also distinguishes them from other *Methanosarcinaceae* but not *Methanothrix* (Tab. S4). These primarily appear to be type II TA systems such as *hicAB*, *relBE*, *mazEF*, *vapBC*, and MNT-HEPN (*143*). The presence of these TA systems may be related to the enrichment in mobile elements, as some type II TA systems may serve to stabilize and maintain horizontally transferred genomic islands or transposons (*89*). However, TA systems have also been associated with other functions relevant to ANME biology. In particular, the *relBE* and *mazEF* systems have been implicated in increased biofilm formation in some bacteria (*89*). They and others may also play a role in the bacterial response to nutrient limitation by inducing reversible growth inhibition (*89*).

ANME and methanogenic *Methanosarcinaceae* also differ in the prevalence of multi-heme cytochrome (MHC) proteins in their genomes. As noted in previous studies, ANME genomes often contain several times more genes for MHCs than related methanogens (*16*, *144*). We observe a similar distribution in our study genomes (Fig. S9). While methanogenic *Methanosarcinaceae* have an average of 24.8 cytochromes and 4.4 MHCs per genome, *Methanocomedenaceae* and *Methanogasteraceae* average 33.4 cytochromes and 15.4 MHCs per genome, and *Methanovorans* average 30.5 cytochromes and 12.1 MHCs (Tab. S5). Interestingly, it appears that there is no significant difference in the abundance of mono-heme cytochromes between these groups (ANOVA: *p >* 0.336). Instead, any difference between the cytochrome complement of these organisms is due to the apparent addition of MHCs to the genome, as *Methanocomedenaceae*, *Methanogasteraceae*, and *Methanovorans* have significantly elevated counts of MHCs compared to these methanogens (ANOVA: *p <* 5.4 *×* 10*^−^*^8^, post-hoc t-tests: ANME-2 vs. methanogens, *p <* 8.2 *×* 10*^−^*^7^; ANME-3 vs. methanogens, *p <* 0.0038). While *Methanovorans* seem to have lower MHC counts than *Methanocomedenaceae* and *Methanogasteraceae* on average, this difference is not significant (post-hoc t-test: *p >* 0.089).

#### 7.2.3 ANME encode distinctive transporter proteins

While nutrient and metabolite transport as a functional class was not found to be enriched in ANME relative to methanogens, ANME do show enrichment in specific sets of transport-related genes. Examining enrichment of functional annotations in the genomes from several databases, we observed *Methanocomendaceae* (ANME-2ab), *Methanogasteraceae* (ANME-2c), and *Methanovorans* (ANME-3) consistently had high enrichment in branched-chain amino acid transport proteins. Functional enrichment analyses also indicated enrichment in up to six other transpor systems, but the predicted substrate of these other transporters was not consistent across annotation databases (file S1). However, several protein families (as defined by the pangenome analysis) annotated as transporters were also enriched in ANME over methanogens (file S1, sheet IDENTITY). These included components of putative phosphate, iron, and multidrug transport systems, each of which was present in at most 2/18 methanogen genomes and at least 17/21 ANME genomes. It has been proposed that ANME partnerships with sulfate reducing bacteria may be shaped by nutrient or vetabolite exchange (*9*), so the increased abundance of certain transporters may reflect this relationship.

ANME also appear to encode a distinctive homolog of the Mrp/Mnh multisubunit sodium-proton antiporter complex. This complex has been suggested to be an important part of energy conversion in diverse archaea (*145*). The existence of a potentially ANME-specific homolog of Mnh is particularly interesting given the observed ANME-specific modification of the energy-conserving sodium transport complex Rnf. The protein family enrichment analysis revealed ANME-specific homologs of MnhBCEFG present in all 11 ANME-2 genomes and about half of the *Methanovorans* genomes (file S1). Further investigation revealed that methanogenic *Methanosarcinaceae* and the other *Methanovorans* genomes encode a different homolog of each of these Mnh subunits (file S2). Within *Methanovorans*, these ANME-specific homologs are present in the entire subclade containing species HMMV2 and SB1 and partially in two Jaco genomes (MAGs Jaco-10860-coassembly and Jaco-11668-bin3). The pattern of presence and absence of these genes in *Methanovorans* genus suggests the ANME-like form was a relatively early acquisition after the genus’ last common ancestor in one *Methanovorans* subclade, with potentially later acquisitions of the same locus in one of the Jaco lineages. While such an early acquisition could indicate some metabolic importance for *Methanovorans*, the presence of the ANME-like Mnh in only half of the genus complicates this interpretation. Additionally, there is not a clear correlation between *Methanovorans* genomes which lack these ANME Mrp/Mnh homologs and other distinctive ANME features.

ANME may also have specific requirements for protein export. *Methanocomendaceae*, *Methanogasteraceae*, and *Methanovorans* genomes contain a highly conserved locus putatively encoding the LolCDE lipoprotein export system. In this system, the homologous LolCE proteins form a dimer in the cell membrane and interact with two copies of the cytoplasmic LolD ATPase to transfer lipoproteins from the membrane to a periplasmic chaperone LolA (*146*). While many of the methanogenic taxa in this study also encode elements of this system, their loci have different gene content from ANME. In ANME, the locus contains *lolCDE* and a conserved unknown protein (Fig. S10). In methanogens, there is variability in the organization of the locus and several genes are frequently absent (Fig. S10). Though it is not present in the locus, at least one gene annotated as *lolA* is encoded by 9/11 ANME-2 genomes and all 10 *Methanovorans* genomes in this study, while only appearing in 11/18 methanogen genomes. Additionally, phylogenies of the genes in the locus clearly separate ANME from methanogens, with *Methanovorans* interspersed among *Methanocomendaceae* and *Methanogasteraceae* (Fig. S11, S12). This suggests *Methanovorans* acquired the locus horizontally from *Methanocomendaceae* and/or *Methanogasteraceae* in multiple separate events. The multiple introductions of these genes may also indicate a viral origin, as observed for *thyX* in *Methanophagales* (ANME-1) (*99*). Oddly, however, the phylogeny of the conserved unknown protein in the locus agrees well with the estimated species phylogeny, clearly separating the ANME clades (Fig. S13).

#### 7.2.4 Potential for methanol- and methylamine-based methanogenesis in *Methanovorans*

Many members of the order *Methanosarcinales* are known to perform methylotrophic methanogenesis (*14*). One such pathway common to many members of the order produces methane from methanol using the proteins MtaABC. While this metabolism is common among methanogenic *Methanosarcinales*, methanotrophic *Methanocomedenaceae* (ANME-2ab) and *Methanogasteraceae* (ANME-2c) lack the pathway (Fig. S1, (*16*)). Only two ANME genomes in this study, both *Methanovorans* (ANME-3) from the Scotian Basin, contain any genes required for methanol-dependent methanogenesis (Fig. S1). In contrast, KEGG annotations indicate that all but one of the methanogen genomes in this study encodes a complete pathway for methanol-dependent methanogenesis (Fig. S1). Each of the two Scotian Basin *Methanovorans* genomes encode a single gene annotated as *mtaA*, but these gene sequences are unlike those from methanogen *mtaABC* loci. The best sequence matches in methanogens to the *Methanovorans mtaA* genes do not share a locus with other methanogenesis genes, suggesting these genes may not truly be associated with methanol-based methanogenesis.

The pathway for methanogenesis from methylamines in *Methanosarcinaceae* consists of the enzymes MtbABC, MtmBC, MttBC, and RamA (*28*). The enzymes MtmB, MtbB, and MttB, which initiate the process for monomethylamine, dimethylamine, and trimethylamine respectively, each require the non-canonical amino acid pyrrolysine (*28*). This further depends on the pyrrolysine biosynthesis genes *pylBCD* as well as a specific tRNA (PylT) and tRNA-aminoacyl synthetase (PylS). As mentioned in the main text, the *Methanovorans* MAGs S4-bin26-refined and S7-bin36-refined (recovered from within or just below the sulfate-methane transition zone) retain all genes required for both pathways. Though their codon usage frequencies suggest that they do not use pyrrolysine, it is possible they may be capable of methylamine-based methanogenesis. In *Methanovorans* MAG HMMV, however, the *pylD* and *pylB* genes in this MAG are truncated. Also, a transposase may have been inserted into the *pylS* gene, though each half of the *pylS* gene is still recognizable through alignment to other *pylS* sequences (file S4).

#### 7.2.5 Conservation in ANME methyl-coenzyme M reductase

The methyl-coenzyme M reductase (Mcr) complex catalyzes the first step in anaerobic methane oxidation (or the last step in methanogenesis). Some recent work suggests that Mcr from *Methanophagales* (ANME-1) may facilitate reversing the methanogenesis pathway in *Methanosarcina* species to perform AOM (*20*), while other work has suggested there are likely minimal structural differences between MCR from methanogens and ANME-2 (*54*). Despite this apparent similarity between methanogen and ANME-2 MCR, which should presumably extend to *Methanovorans* (ANME-3) as well, we observed what appears to be a phylogenetic signal indicating some distinction exists at the sequence level between MCR from these groups (Fig. 3).

As discussed in the main text, the protein sequence alignments for each MCR subunit do not show insertions or deletions that distinguish ANME-like Mcr from methanogen-like Mcr. However, we observe higher rates of ANME-characteristic conserved residues (70% identity) in MCR than in phylogenetic marker genes (Tab. S3). If we slightly relax the identity threshold to include less highly conserved (60% identity) but still ANME-characteristic residues, this difference becomes even more striking (13.1 per kAA in MCR, 4.9 per kAA in marker genes). This difference in rates at the 60% identity threshold is significant based on 95% confidence intervals (Tab. S3).

It is difficult to interpret the effect that these conserved residues may have on the function or stability of ANME Mcr. In some of these sites, ANME and methanogens seem to have amino acid residues with similar properties but different side chain sizes. In other sites, ANME may have replaced polar or charged residues with hydrophobic residues, or vice versa (files S5, S6, S7, S8, S9, S10). Interestingly, in two of these sites ANME appear to have lost a proline which is highly conserved in methanogens. Structural predictions of ANME Mcr subunits suggest that the sites where ANME have conserved charged residues often sit near the interface between McrA and other subunits in the Mcr complex, but do not sit near the active site (supp. file S15). One potential explanation for this observation is that these residues are important for posttranslational modifications of MCR. In *Methanophagales* (ANME-1), the Mcr enzyme complex undergoes many post-translational modifications which distinguish it from methanogenic Mcr structures (*53*). However, Shima *et al.* note that several of these are unlikely to be present in *Methanocomedenaceae* (ANME-2ab), *Methanogasteraceae* (ANME-2c), or *Methanovorans* (ANME-3), complicating their relationship to AOM. The high prevalence of diverse and taxonomically heterogeneous post-translational modifications to ANME Mcr, often far from the active site, has also been noted in proteomic data sets, suggesting these modifications may be responses to environmental variation rather than adaptations to enhance AOM (*147*).

#### 7.2.6 Conservation in the ANME RNF locus

In *Methanosarcina acetivorans* and other methanogenic *Methanosarcinaceae*, the Rhodobacter nitrogen fixation (RNF) locus consists of the six genes *rnfABCDEG*, a multiheme cytochrome *mmcA*, and one additional conserved gene (*148*). As has been previously noted, the ANME RNF locus additionally encodes a cytochrome b protein and two short transmembrane hairpin proteins which are not present in other *Methanosarcinaceae* (*16*). The MmcA and cytochrome b proteins are notable as they may modify electron flow through the complex, and recent studies have suggested that MmcA may facilitate intracellular and extracellular electron transport (*63*, *62*).

As mentioned in the main text, the RnfC protein in ANME has an N-terminal domain feature which appears to be distinctive. Based on sequence alignments of the *rnfC* gene, this domain represents a modification to the existing N-terminal sequence rather than an extension of the protein. Structural predictions of RnfC suggest that this N-terminal region in *Methanovorans* is more similar to ANME-2 RnfC than RnfC from *Methanococcoides*, further suggesting that *Methanovorans* RNF has undergone convergent modifications toward other ANME. In fact, this N-terminal region accounts for much of the difference in the root-mean-squared distance (RMSD) between the structures of the three groups. In ANME the region appears to have fewer protrusions and a small indentation (potentially a binding pocket) (Fig. S4). While the surface potential varies in each organism, in both ANME this N-terminal region has a more negative potential than the rest of the protein, while in *Methanococcoides* the region has a more positive potential. Based on a topological prediction of the Rnf complex in *Vibrio cholerae*, it appears that this altered region of RnfC would sit very close to the cytoplasmic side of the cell membrane (*59*). It seems possible that this domain could facilitate association of the additional ANME-specific components to the rest of the complex.

The main text also mentions modifications to the MmcA protein sequence, which will be described here in more detail (see file S11). In a protein sequence alignment of ANME and methanogen *mmcA*, *Methanovorans* have deletions (relative to the methanogen sequences) at a similar position to a gap in the *Methanocomedenaceae* sequences. Elsewhere in the alignment there is a short domain present only in *Methanovorans* and *Methanolobus* species. Most intriguingly, *Methanovorans* and *Methanocomedenaceae mmcA* genes seem to share a similar heme binding environment which is distinct from methanogens. In the methanogen genomes in this study, *mmcA* has 7 or 8 heme binding motifs, including one CXXXCH motif and one CX_4_CH motif. *Methanovorans mmcA* has 8 heme-binding motifs and in *Methanocomedenaceae* the gene has 9 heme-binding motifs, including two at positions which are not shared with either methanogens or *Methanovorans*. Oddly, all *Methanosarcinaceae* genomes have a conserved YXXYH motif at one of the two *Methanocomedenaceae*-specific positions and a fairly conserved [FY]XXAN motif at the other. The potential for these motifs to bind heme b instead of heme c is worthy of further investigation. Also, in the position where methanogens have the unusual CX_4_CH motif, all *Methanocomedenaceae* and *Methanovorans* genomes instead have a highly conserved CIDCH heme-binding motif. In bacteria, the conversion of a heme binding motif from CX_4_CH to CXXCH significantly alters the conformation of the bound heme, and therefore may affect the redox potential at which it is poised (*149*, *150*). This highly conserved binding motif in ANME suggests a very specific redox potential may be required at this heme in ANME metabolism. Given the distinct ancestral background of *Methanocomedenaceae*, *Methanogasteraceae*, and *Methanovorans*, this suggests a similar selective pressure could be shaping these proteins towards similar properties.

#### 7.2.7 Novel large ANME multiheme cytochromes in *Methanovorans*

Based on their distribution within *Methanovorans*, it is likely that the large ANME MHC proteins were acquired relatively early after the divergence of the clade. One deeply branching *Methanovorans* subclade contains the MHC-A locus, and another contains the MHC-B/C locus or genes derived from it via fusion. Though *Methanovorans* MAG HMMV and *Methanovorans* MAG HMMV2 were collected from the same site, they sit in opposite subclades and have different MHC loci. Interestingly, recent studies of *Methanosarcina* have indicated that the MmcA protein, in conjunction with the Rnf complex, may allow direct electron exchange with the environment (*61*). This suggests that these large MHC proteins are not fundamentally essential to ANME metabolism, though they may offer a significant advantage.

In most *Methanocomendaceae* and *Methanogasteraceae*, MHC-A is a protein with roughly 1200 amino acids, one or two transmembrane helices, and an S-layer domain. In *Methanovorans* the gene has a similar architecture of transmembrane helices and S-layer domains, but codes for a protein with roughly 2400 amino acids. *Methanovorans* MHC-A has relatively low sequence identity and locus synteny with most other *Methanocomendaceae* or *Methanogasteraceae* in this study, with the exception of one *Methanocomedens* (ANME-2a) single aggregate genome with an unusually large MHC-A. BLASTP searches against the NCBI nr database show no sequences outside of ANME with decent coverage and identity. Many sequences in other publicly available *Methanocomendaceae* or *Methanogasteraceae* assemblies show 50–60% sequence identity to regions of the *Methanovorans* genes but are significantly shorter, much like most *Methanocomendaceae* or *Methanogasteraceae* MHC-A sequences in this study. Several studies have noted that *Methanoperedens* genomes often include unusually large and diverse MHC sequences e.g. (*144*). While there are several public sequences from *Methanoperedens* species with similar lengths to the *Methanovorans* MHC-A genes, these typically have only 30–40% sequence identity to *Methanovorans*. This is enough similarity to indicate a potential common origin, but it does not appear that ANME-3 acquired their large MHCs from *Methanoperedens* directly.

In *Methanocomedenaceae*, MHC-B and -C co-occur in a locus together with several components of cytochrome c maturation machinery. One *Methanovorans* genome has a locus encoding these proteins with good sequence identity and high synteny with *Methanocomedens* genomes, suggesting it was acquired by horizontal gene transfer from *Methanocomedens* (Fig. 5). However, two *Methanovorans* genomes contain a large MHC protein (MHC-D) which appears to be a fusion of MHC-B and MHC-C. While MHC-B is typically 2200 amino acids and MHC-C is 2400 amino acids, the fusion MHC-D is about 3200 amino acids long. The N-terminal region of MHC-D aligns well with the N-terminal region of MHC-C, and the C-terminal region aligns well with the C-terminal region of MHC-B (files S12, S13). This leads to an overall domain structure of MHC-D with a C-terminal transmembrane helix and S-layer domain and an N-terminal pectin lyase-like domain, with heme binding sites throughout. Given the fact that MHC-B and MHC-C share a locus and a fusion of the two exists, it seems likely that these two proteins may interact and function together, with MHC-B serving as a conduit for electrons to leave the ANME cell and interacting with MHC-C through its peptidase M6-like domain, and MHC-C potentially interacting with the cell surface of the sulfate-reducing partner through its pectin lyase domain. In this case, MHC-D may not serve a different function, but may trade (for example) easier maturation and export for shorter interaction distance. Interestingly, one *Methanovorans* genome contains both the MHC-D locus and the MHC-B/MHC-C locus. However, gene phylogeny indicates that the MHC-D gene is more similar to MHC-D from other *Methanovorans* than to the unfused MHC-B and MHC-C within the same genome (Fig. S14). This suggests that this *Methanovorans* did not undergo duplication and fusion of the locus, but rather independently acquired both the fused and unfused versions by horizontal transfer.

#### 7.2.8 Contractile injection systems in *Methanovorans*

Phage-like protein translocation systems (PLTS) are structurally similar to other phage-derived contractile injection systems (CIS) such as type VI secretion systems and R-type pyocins but share only distant evolutionary roots (*71*, *72*). PLTS are broadly distributed across the old *Euryarchaeota* phylum and several bacterial phyla, with several core structural components as well as variable supplementary genes which may modify the systems for more specialized functions (*151*). Much of the variability in function can likely be attributed to the variety of effector proteins which can be delivered by these systems. Unfortunately, while strides have been made allowing identification of potential toxin effectors in diverse bacterial taxa, archaeal effectors have proved more difficult to identify (*151*).

Euryarchaeal CIS are not monophyletic, indicating at least two primary acquisitions of the cluster from bacteria (*151*). Interestingly, *Methanocomedenaceae* (ANME-2ab) CIS do not all cluster together. While it seems like CIS in *Methanovorans*, *Methanogaster*, and some *Methanocomedenaceae* stem from a single primary acquisition in *Euryarchaeota*, CIS in other *Methanocomedenaceae* and methanogenic archaea may stem from a separate acquisition event. This suggests that there may be secondary horizontal transfer of CIS within the archaea leading to the acquisition of the system by *Methanovorans*

*Methanovorans* CIS loci universally lack an ATPase gene typically associated with PLTS, in contrast to other *Methanosarcinaceae* and other ANME loci. In several cases, the synteny of the surrounding genes is the same between *Methanocomedenaceae*, *Methanogasteraceae*, and *Methanovorans*, suggesting that this loss is specific and not due to a larger-scale rearrangement of genes. This is particularly interesting if, as suggested above, each of these *Methanovorans* CIS were acquired independently and thus this gene was lost independently multiple times. In type VI secretion systems, this ATPase is thought to be involved in disassembly of the tube after contraction (*152*, *76*). However, it was not necessary for assembly of complete particles in an extracellular CIS (*35*).

#### 7.2.9 Functionality of nitrogenase in ANME

ANME nitrogenase genes tend to be distinct from most other bacterial or archaeal nitrogenases (*70*, *153*). Using structural prediction software revealed that one distinguishing feature may be a more solvent-exposed active site in ANME nitrogenase, potentially altering its activity or specificity (Fig. S8). It is worth noting that nitrogenase is known to have functions other than nitrogen fixation, including reduction of carbon monoxide and other small molecules (*154*). Another notable feature is that all of the ANME *nif* loci lack *nifEN*, and these genes were not found elsewhere in any ANME genomes in this study. The genes *nifEN* are considered essential for the proper synthesis and incorporation of the P-cluster and MoFe cofactor into nitrogenase enzymes (*155*). Despite this, previous work has shown that ANME-2 actively fix nitrogen both in lab incubations and *in situ* on the ocean floor (*70*, *86*). The *nifD* sequences found in the genomes in this study are close matches to methane seep sequences from those studies (up to 94% identical), suggesting they are likely to be functional.

It is unclear how the ANME may compensate for the absence of the NifEN subunits. However, as *nifDK* and *nifEN* are close homologs, it is possible that *nifDK* may be able to take on the role of *nifEN* in ANME. It appears that when using the alternative nitrogenase Anf, *nifEN* are not required for nitrogen fixation (*156*). There is also some evidence for other organisms which appear to lack *nifEN*. For example, several *Roseiflexus* and *Aquificales* isolates encode *nif* genes but lack *nifE* and/or *nifN*, though nitrogenase activity has not been reported for them (*157*). A *Methanocaldococcus* strain was recently reported to fix nitrogen despite not encoding *nifN* (*158*). More intriguingly, nitrogen fixation has been reported in *Endomicrobium proavitum* which encodes a group IV nitrogenase but does not encode *nifEN* (*159*), suggesting that these genes may not be required in all cases for nitrogenase function.

#### 7.2.10 *Methanovorans* encode ANME-like and methanogen-like Hdr

The heterodisulfide reductase enzymes are found in both methanogens and ANME and couple the oxidation of the coenzyme M-coenzyme B heterodisulfide with the reduction of methanophenazine, ferredoxin, or cofactor F_420_ (*160*). The HdrDE complex is membrane-bound and the HdrABC complex is soluble, but organisms may encode both versions of the complex. ANME encode diverse Hdr enzymes with multiple homologs of HdrA implicated in electron bifurcation and/or confurcation, which has been suggested to be an important novelty in ANME metabolism (*16*). Our protein family enrichment analysis revealed several *hdr* genes are heavily enriched in ANME over methanogenic *Methanosarcinaceae*, some of which were exclusive to ANME (file S1). This was true for both the soluble and membrane-bound Hdr complexes. Further examination also revealed *hdr* genes exclusive to *Methanosarcinaceae* (file S2). Intriguingly, some *Methanovorans* genomes exclusively encode *hdr* genes specific to ANME, others encode exclusively encode *hdr* genes specific to *Methanosarcinaceae*, and some encode at least one gene from each of the prior categories (sometimes in the same locus). This supports the hypothesis that the evolutionary transition of *Methanovorans* toward methanotrophy is ongoing. Also, regardless of their origin, ANME-like *hdr* is not a conserved trait in *Methanovorans* and thus its acquisition is unlikely to be a precipitating event in their evolution towards methanotrophy.

### 7.3 Supplementary figures and tables

**Table S1.**
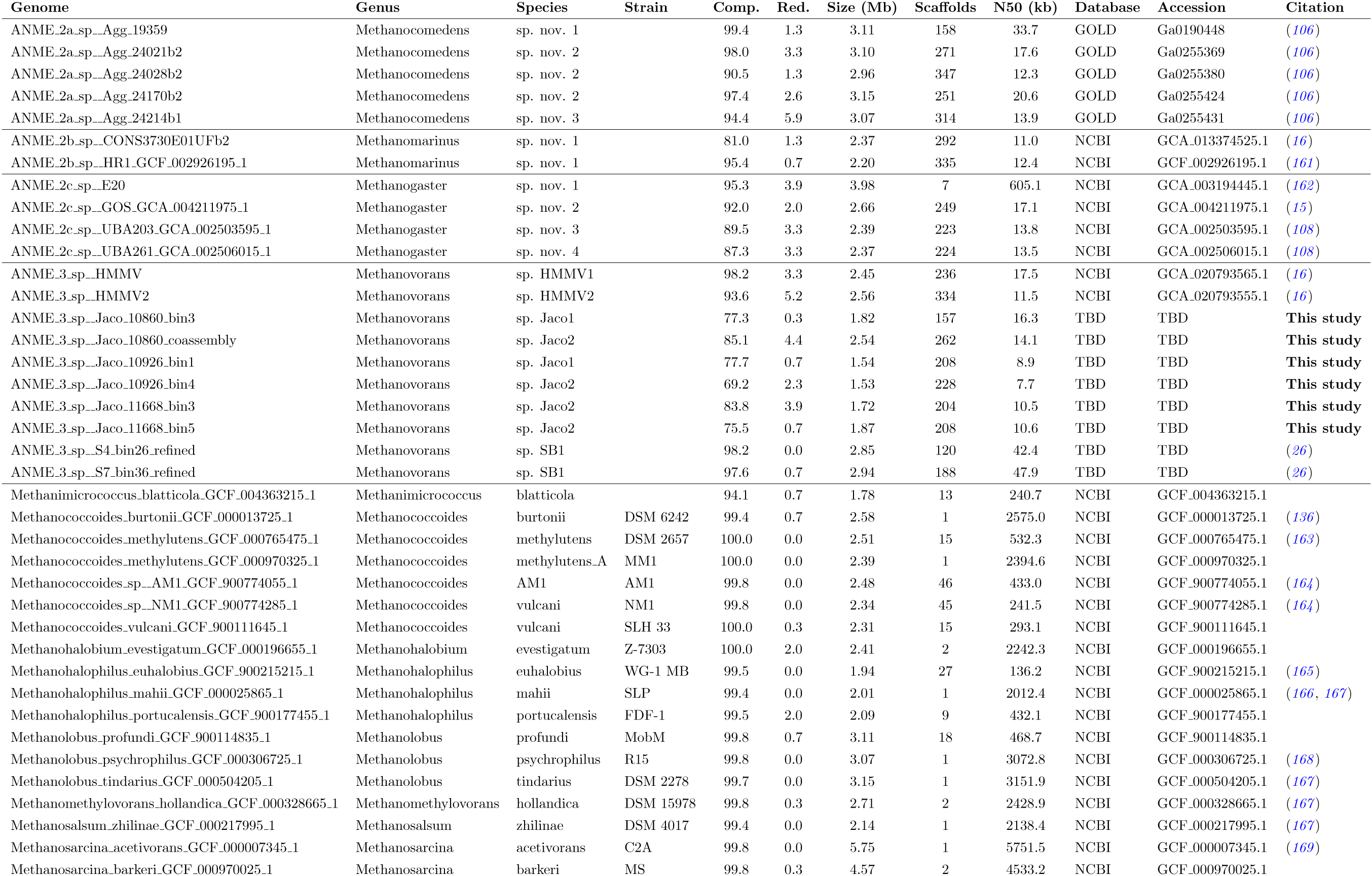
Summary of genome statistics, taxonomy, accessions, and citations for genomes used in this study. Taxonomy follows the GTDB standard. Genus names for ANME follow Chadwick *et al.* (*16*). Estimated completeness (Comp.) and redundancy (Red.) are reported as percent.

**Table S2.** Stop codon frequency in ANME and methanogen genomes. Pyrrolysine-utilizing methanogens have very low frequencies of the TAG stop codon, which is also used to encode Pyl. In ANME-2 and *Methanothrix*, which do not use pyrrolysine, the TAG stop codon occurs at higher rates. ANME-3, including those genomes which posess genes for pyrrolysine and methylamine metabolism, have similar TAG codon frequencies to ANME-2. (*) Codon usage for *Methanosaeta thermophila* from Alkalaeva *et al.* (*30*).

| Genome name | Stop codon frequency |  |  | Gene content |  |
| --- | --- | --- | --- | --- | --- |
|  | TAA | TAG | TGA | Pyl synthesis & expression | Methylamine methanogenesis |
| Methanosaeta harundinacea 57_489 | 17.5 | 20.1 | 62.4 | - | - |
| Methanosaeta harundinacea UBA475 | 13.5 | 21.7 | 64.8 | - | - |
| Methanosaeta harundinacea 6Ac | 11.5 | 21.2 | 67.3 | - | - |
| Methanosaeta sp. PtaB_Bin087 | 11.4 | 19.9 | 68.7 | - | - |
| Methanosaeta sp. PtaU1_Bin060 | 19.2 | 24.9 | 55.9 | - | - |
| Methanosaeta sp. PtaU1_Bin112 | 25.9 | 25.6 | 48.5 | - | - |
| Methanosaeta sp. UBA114 | 29.4 | 26.6 | 43.9 | - | - |
| Methanosaeta sp. UBA204 | 22.5 | 23.7 | 53.8 | - | - |
| Methanotherix soehngenii | 25.7 | 25.9 | 48.4 | - | - |
| Methanotherix sp. fen_7 | 29.4 | 25.4 | 45.2 | - | - |
| Methanotherix thermoacetophila | 14.6 | 18.9 | 66.5 | - | - |
| Methanosaeta thermophila * | 12.8 | 19.1 | 68.1 | - | - |
| ANME-2a MAG Agg-19359 | 39.6 | 18.6 | 41.9 | - | - |
| ANME-2a MAG Agg-24021b2 | 37.3 | 19.4 | 43.3 | - | - |
| ANME-2a MAG Agg-24028b2 | 38.3 | 19.1 | 42.6 | - | - |
| ANME-2a MAG Agg-24170b2 | 38.5 | 19.2 | 42.3 | - | - |
| ANME-2a MAG Agg-24214b1 | 39.5 | 19.3 | 41.1 | - | - |
| ANME-2b MAG CONS3730E01UFb2 | 46.8 | 17.9 | 35.3 | - | - |
| ANME-2b MAG HR1 | 45.5 | 18.9 | 35.6 | - | - |
| ANME-2c MAG E20 | 28.1 | 17.8 | 54.1 | - | - |
| ANME-2c MAG GOS | 30.0 | 15.6 | 54.4 | - | - |
| ANME-2c MAG UBA203 | 24.8 | 16.0 | 59.2 | - | - |
| ANME-2c MAG UBA261 | 20.9 | 17.1 | 62.1 | - | - |
| ANME-3 MAG Jaco-10860-bin3 | 42.7 | 18.6 | 38.7 | - | - |
| ANME-3 MAG Jaco-10860-coassembly | 43.2 | 18.6 | 38.2 | - | - |
| ANME-3 MAG Jaco-10926-bin1 | 39.7 | 17.4 | 42.8 | - | - |
| ANME-3 MAG Jaco-10926-bin4 | 41.5 | 17.4 | 41.1 | - | - |
| ANME-3 MAG Jaco-11668-bin3 | 41.4 | 16.5 | 42.2 | - | - |
| ANME-3 MAG Jaco-11668-bin5 | 41.1 | 17.9 | 41.0 | - | - |
| ANME-3 MAG HMMV2 | 41.0 | 17.1 | 42.0 | - | - |
| ANME-3 MAG HMMV | 38.5 | 17.6 | 43.8 | partial | partial |
| ANME-3 MAG S4-bin26-refined | 38.4 | 16.5 | 45.1 | + | + |
| ANME-3 MAG S7-bin36-refined | 39.0 | 17.5 | 43.5 | + | + |
| Methanococcoides burtonii | 45.8 | 3.9 | 50.4 | + | + |
| Methanococcoides methylutens | 46.7 | 2.0 | 51.2 | + | + |
| Methanococcoides methylutens_A | 43.4 | 1.8 | 54.7 | + | + |
| Methanococcoides sp. AM1 | 46.3 | 2.1 | 51.6 | + | + |
| Methanococcoides sp. NM1 | 46.8 | 2.6 | 50.6 | + | + |
| Methanococcoides vulcani | 46.9 | 2.3 | 50.7 | + | + |
| Methanohalophilus euhalobius | 46.8 | 2.5 | 50.7 | + | + |
| Methanohalophilus mahii | 47.5 | 2.0 | 50.5 | + | + |
| Methanohalophilus portucalensis | 48.6 | 1.8 | 49.6 | + | + |
| Methanolobus profundus | 39.8 | 11.1 | 49.1 | + | + |
| Methanolobus psychrophilus | 40.4 | 12.2 | 47.3 | + | + |
| Methanolobus tindarius | 49.6 | 9.1 | 41.3 | + | + |
| Methanomethylovorans hollandica | 42.5 | 9.7 | 47.7 | + | + |
| Methanohalobium evestigatum | 66.7 | 2.2 | 31.1 | + | + |
| Methanosalsum zhilinae | 46.4 | 5.7 | 47.9 | + | + |
| Methanimicrococcus blatticola | 61.5 | 2.0 | 36.5 | + | + |
| Methanosarcina acetivorans | 48.5 | 5.7 | 45.8 | + | + |
| Methanosarcina barkeri | 57.2 | 5.7 | 37.1 | + | + |

**Table S3.** Comparison of conserved residue frequencies between ANME-2 and ANME-3 in Mcr genes and in conserved phylogenetic marker genes. Mcr genes have a much higher rate of highly conserved residues between ANME-2 and ANME-3 than phylogenetic marker genes. This difference in rates becomes more extreme when the conservation threshold is lowered from 70% (top table) to 60% (bottom table).

| Sequence(s) | Conserved residues (70%) | Length (AA) | Conserved residues per kAA | 95% CI |
| --- | --- | --- | --- | --- |
| mcrA | 4 | 584 | 6.8 | (1.9, 17.5) |
| mcrB | 1 | 518 | 1.9 | (0.0, 10.8) |
| mcrG | 3 | 267 | 11.2 | (2.3, 32.8) |
| All MCR | 8 | 1369 | 5.8 | (2.5, 11.5) |
| SCGs | 39 | 14625 | 2.7 | (1.9, 3.6) |
| Archaea76 markers | 4 | 2109 | 1.9 | (0.5, 4.9) |
| All markers | 43 | 16734 | 2.6 | (1.9, 3.5) |

| Sequence(s) | Conserved residues (60%) | Length (AA) | Conserved residues per kAA | 95% CI |
| --- | --- | --- | --- | --- |
| mcrA | 7 | 584 | 12.0 | (4.8, 24.7) |
| mcrB | 6 | 518 | 11.6 | (4.3, 25.2) |
| mcrG | 5 | 267 | 18.7 | (6.1, 43.7) |
| All MCR | 18 | 1369 | 13.1 | (7.8, 20.8) |
| SCGs | 75 | 14625 | 5.1 | (4.0, 6.4) |
| Archaea76 markers | 7 | 2109 | 3.3 | (1.3, 6.8) |
| All markers | 82 | 16734 | 4.9 | (3.9, 6.1) |

**Table S4.** Median abundance of toxin-antitoxin (TA) systems within the COG category “Defense systems” (V). In *Methanovorans* TA systems are a much higher proportion of the annotated defense systems than in methanogenic *Methanosarcinaceae*, but more similar in abundance to *Methanothrix*.

| Grouping | Median TA abundance |
| --- | --- |
| ANME2 | 44% |
| *Methanovorans* (ANME3) | 25% |
| Methanogenic *Methanosarcinaceae* | 8% |
| *Methanotherix* | 32% |

**Table S5.** Average counts of putative cytochrome c proteins in ANME-2, ANME-3, and related methanogens. ANME appear to have more cytochrome c proteins than methanogens. When single-heme and multi-heme proteins are separated, the three groups have similar amounts of single-heme proteins per genome, but ANME-2 and ANME-3 have higher numbers of multi-heme cytochrome proteins than related methanogens.

| Group | All cyt c | (stdev) | Single-heme cyt c | (stdev) | MHCs | (stdev) |
| --- | --- | --- | --- | --- | --- | --- |
| ANME-2 | 33.4 | (4.7) | 18.0 | (3.6) | 15.4 | (3.9) |
| ANME-3 | 30.5 | (11.0) | 18.4 | (5.4) | 12.1 | (6.3) |
| Methanogen | 24.8 | (5.3) | 20.4 | (4.7) | 4.4 | (2.1) |

**Figure S1.**
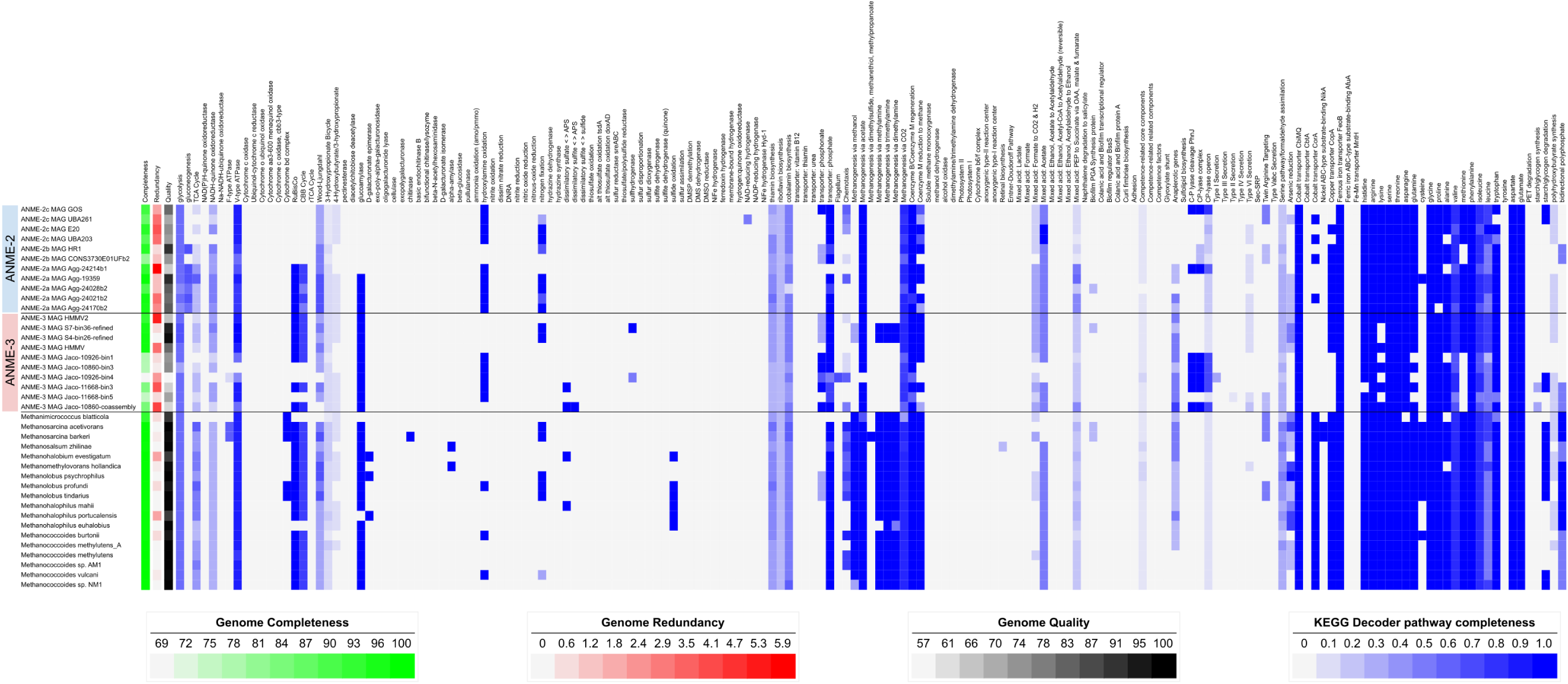
Completeness of metabolic pathways in ANME-2, ANME-3, and related methanogens.

**Figure S2.**
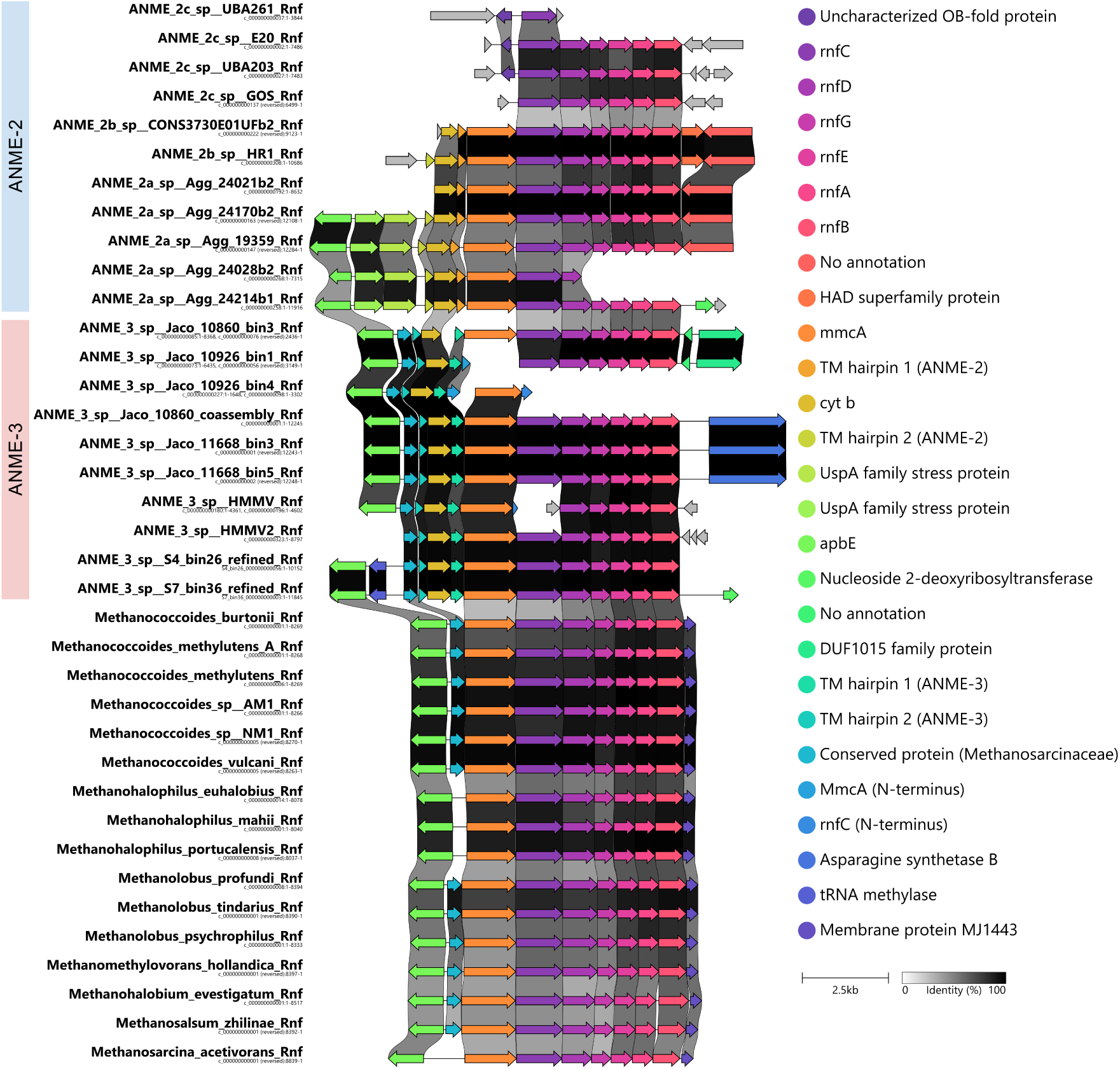
Synteny diagram of the Rnf locus in all study genomes. Note that some ANME-3 have only partial RnfC or MmcA genes due to contig boundaries, but the rest of the locus in those genomes is highly similar to other ANME-3.

**Figure S3.**
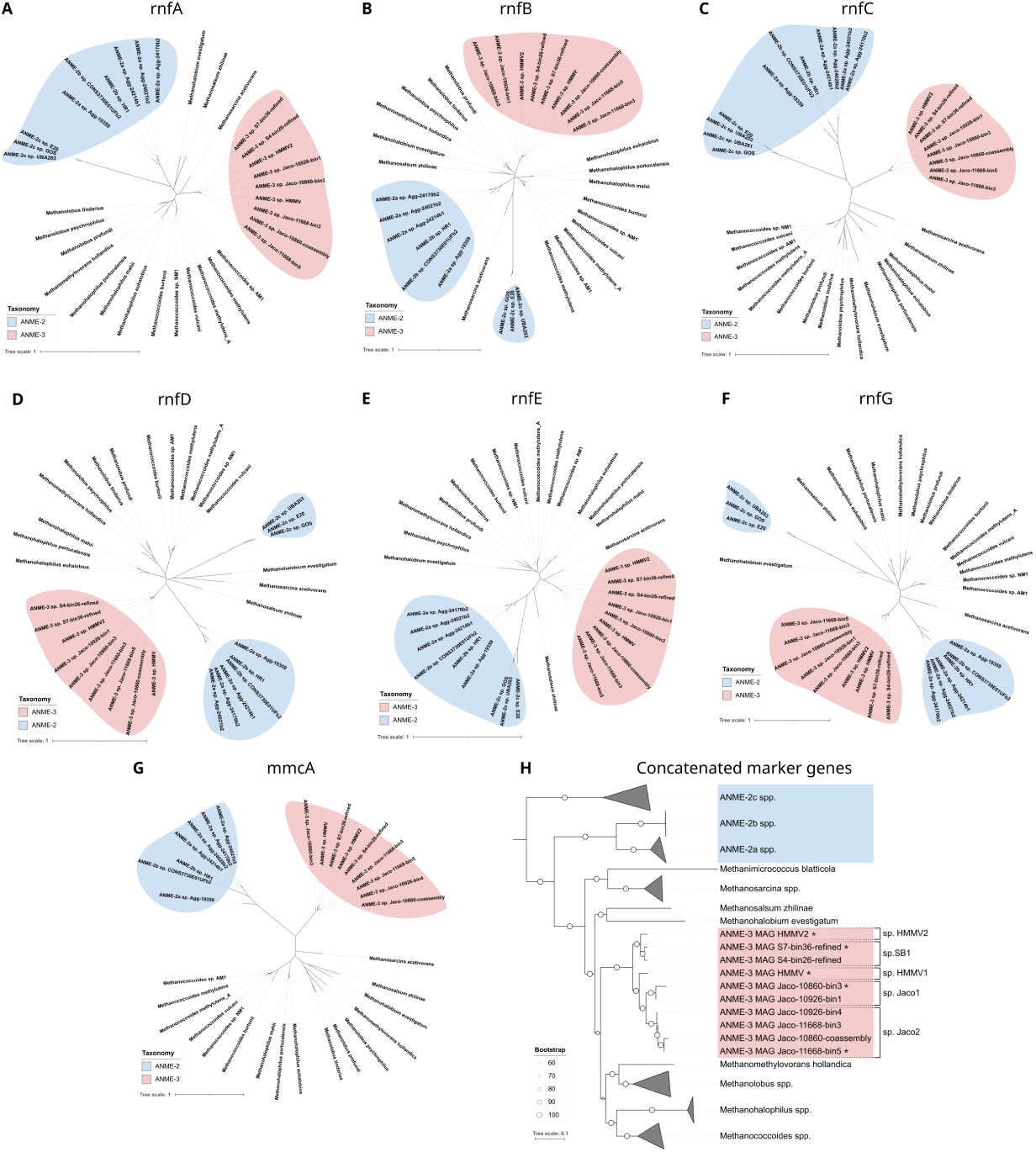
Gene phylogenies of components of the RNF complex. Some components (A, B, E) appear to concord with the estimated species phylogeny (H) while others place ANME-3 closer to ANME-2 (C, D, F, G).

**Figure S4.**
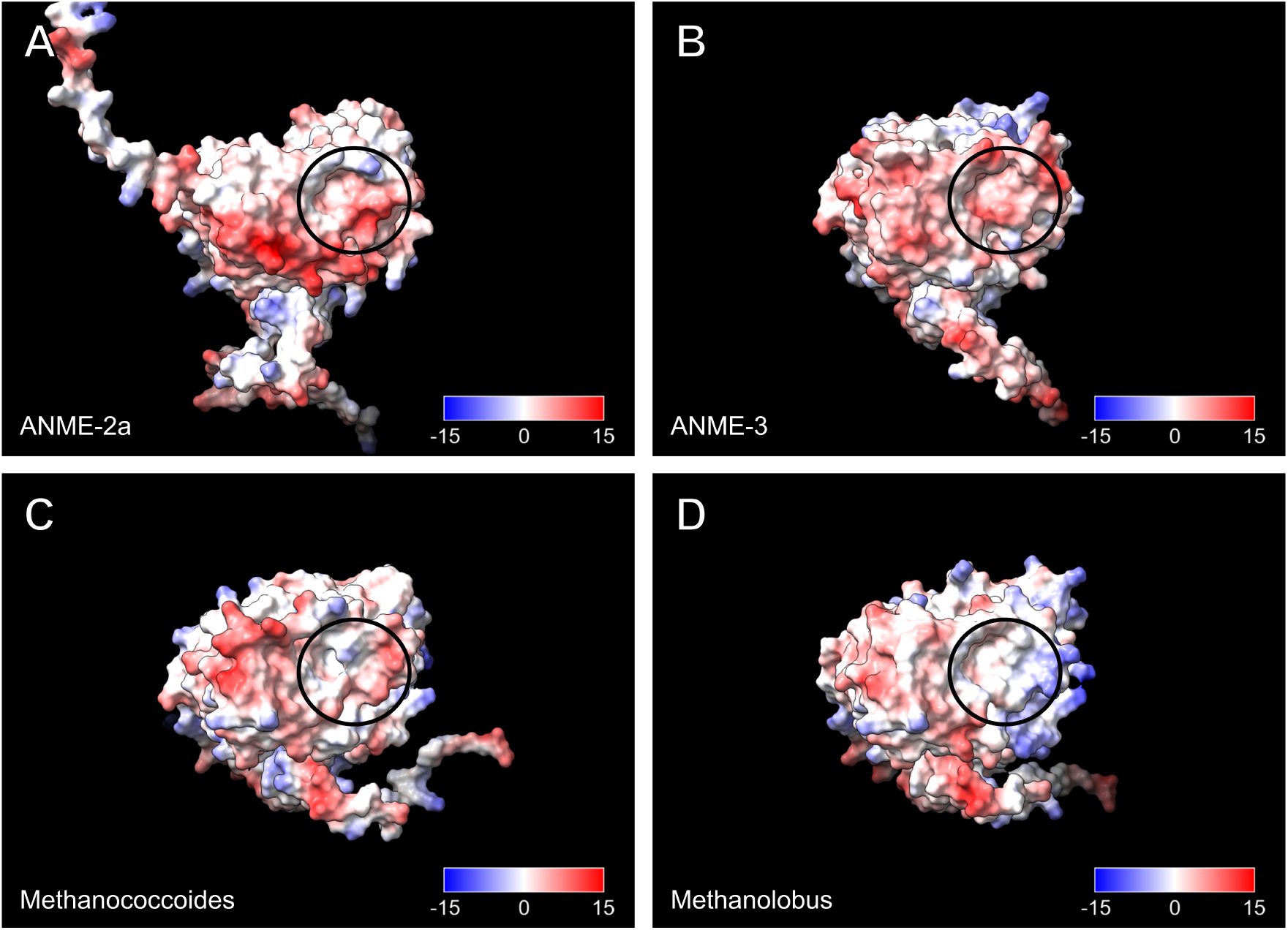
Comparison of predicted structures or RnfC from (**A**) *Methanocomedens* (ANME-2a), (**B**) *Methanovorans* (ANME-3), (**C**) *Methanococcoides*, and (**D**) *Methanolobus*. The ANME structures have a small binding pocket (circled) near the N-terminal region (to the left of the circles). These two regions have more positive surface potential in ANME than in the methanogen representatives.

**Figure S5.**
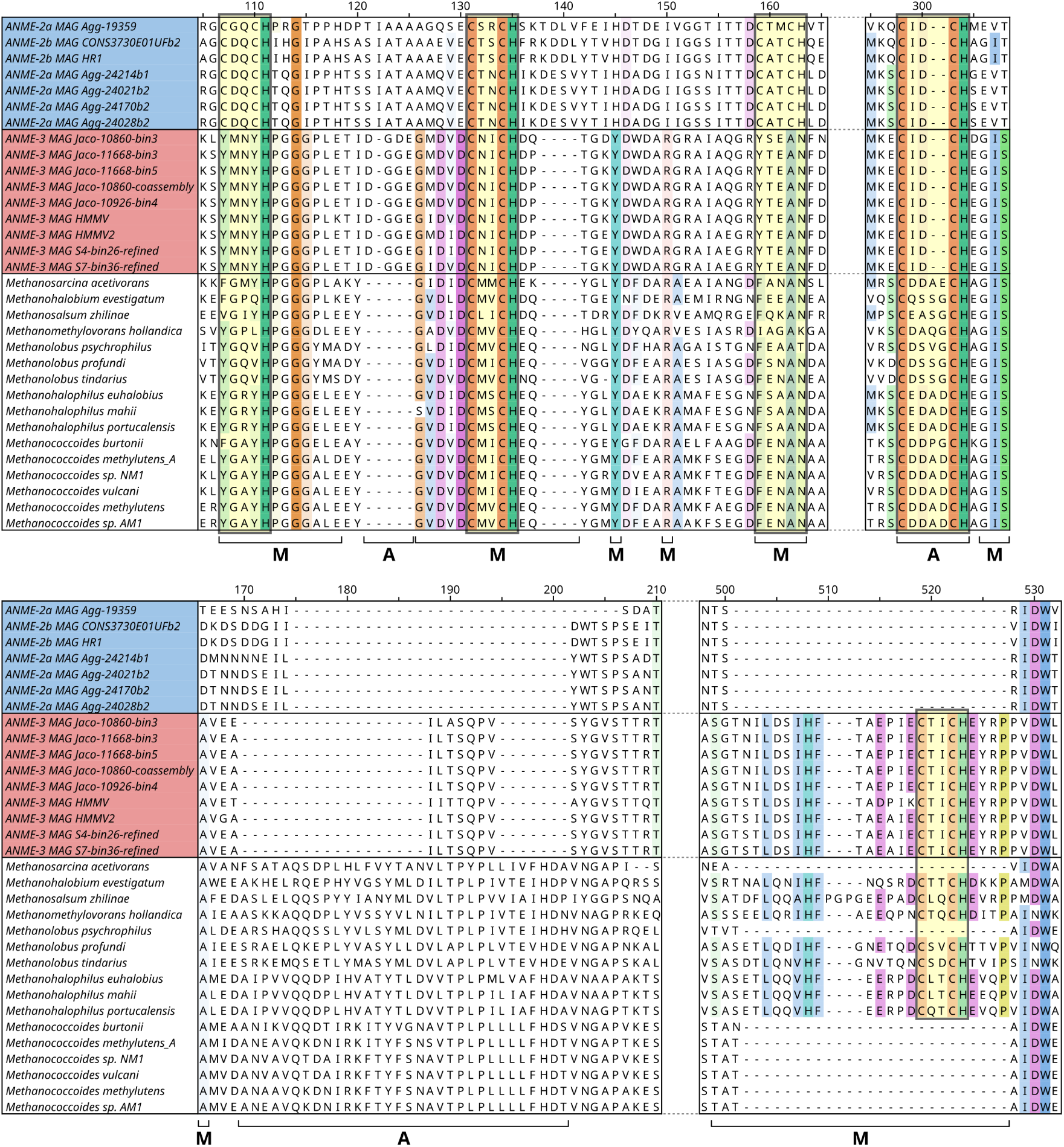
Comparison of insertions, deletions, and heme binding sites in multi-heme cytochrome MmcA. Heme binding motifs are highlighted in in yellow. Brackets below the alignment indicate regions where *Methanovorans* (ANME-3) are similar to *Methanocomedenaceae* (ANME-2ab) (A) or methanogens (M). While ANME share high identity in some regions (e.g. position 298-304), *Methanovorans* generally have more similarity to methanogenic taxa (e.g. 107-118, 126-130, 138-141, 499-527). When there are indels with similar positions and lengths in *Methanocomedenaceae* and *Methanovorans* (e.g. 121-125, 170-201), the sequence and/or surrounding regions typically have low or no conservation.

**Figure S6.**
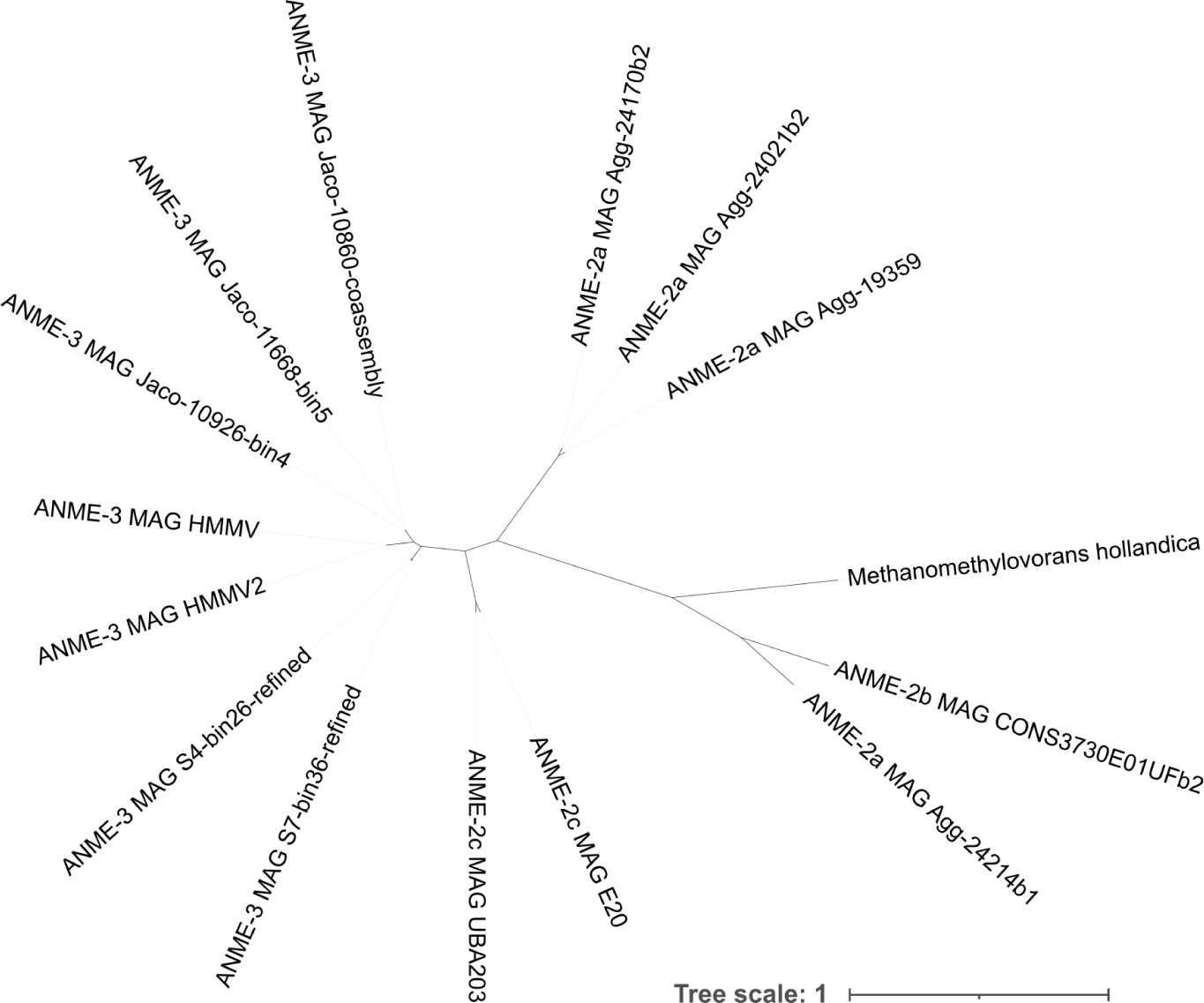
Concatenated gene phylogeny of common core phage-like protein translocation system (PLTS) genes. *Methanovorans* (ANME-3) cluster most closely with *Methanogaster* (ANME-2c). Within *Methanovorans*, taxa cluster most closely with others from the same geographic location. Some *Methanocomedens* (ANME-2a) and *Methanomarinus* (ANME-2b) loci appear to be relatively distantly related to other ANME PLTS loci, including other *Methanocomedens* loci. Genes included and/or excluded from this concatenation are listed in file S14.

**Figure S7.**
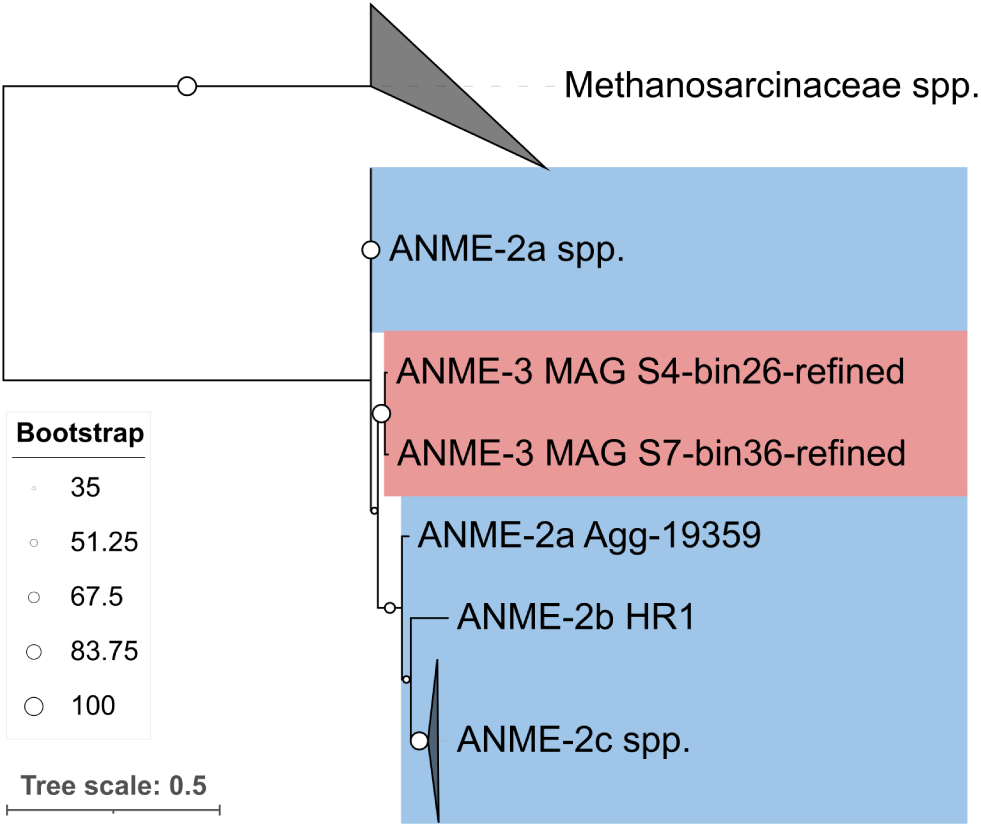
Gene phylogeny of nifD. The gene phylogeny of nitrogenase subunit nifD places ANME-3 nitrogenase among the ANME-2a nitrogenase, distant from the nitrogenase genes in their methanogen relatives.

**Figure S8.**
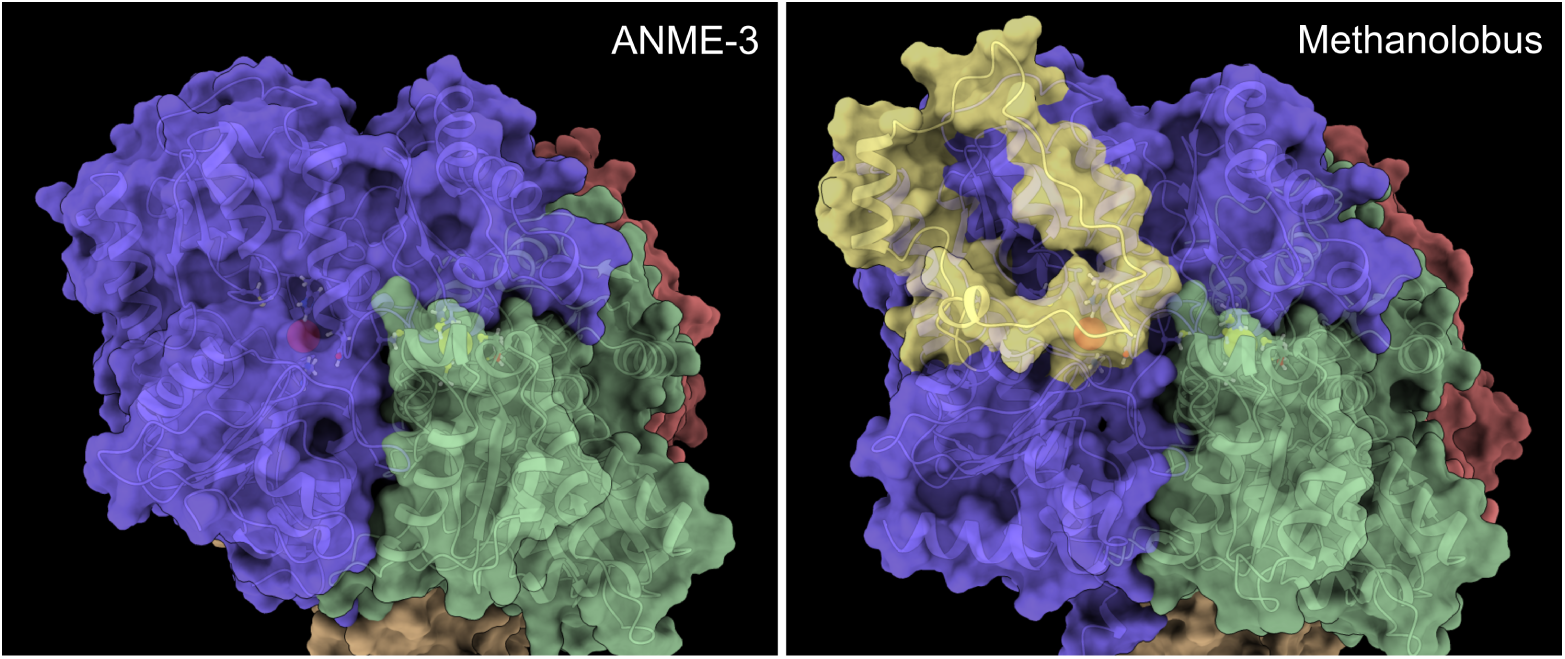
Predicted structures of representative ANME and methanogen NifDK complexes at the FeMo cofactor. NifD (purple, red) and NifK (green, orange) associate to form a tetramer complex. The P-cluster (yellow sphere) bridges the interface between NifD and NifK, while the FeMo cofactor (red sphere) is fully contained within NifD. ANME NifD lacks a surface loop present in methanogens (highlighted in yellow), which may leave the FeMo cofactor more exposed.

**Figure S9.**
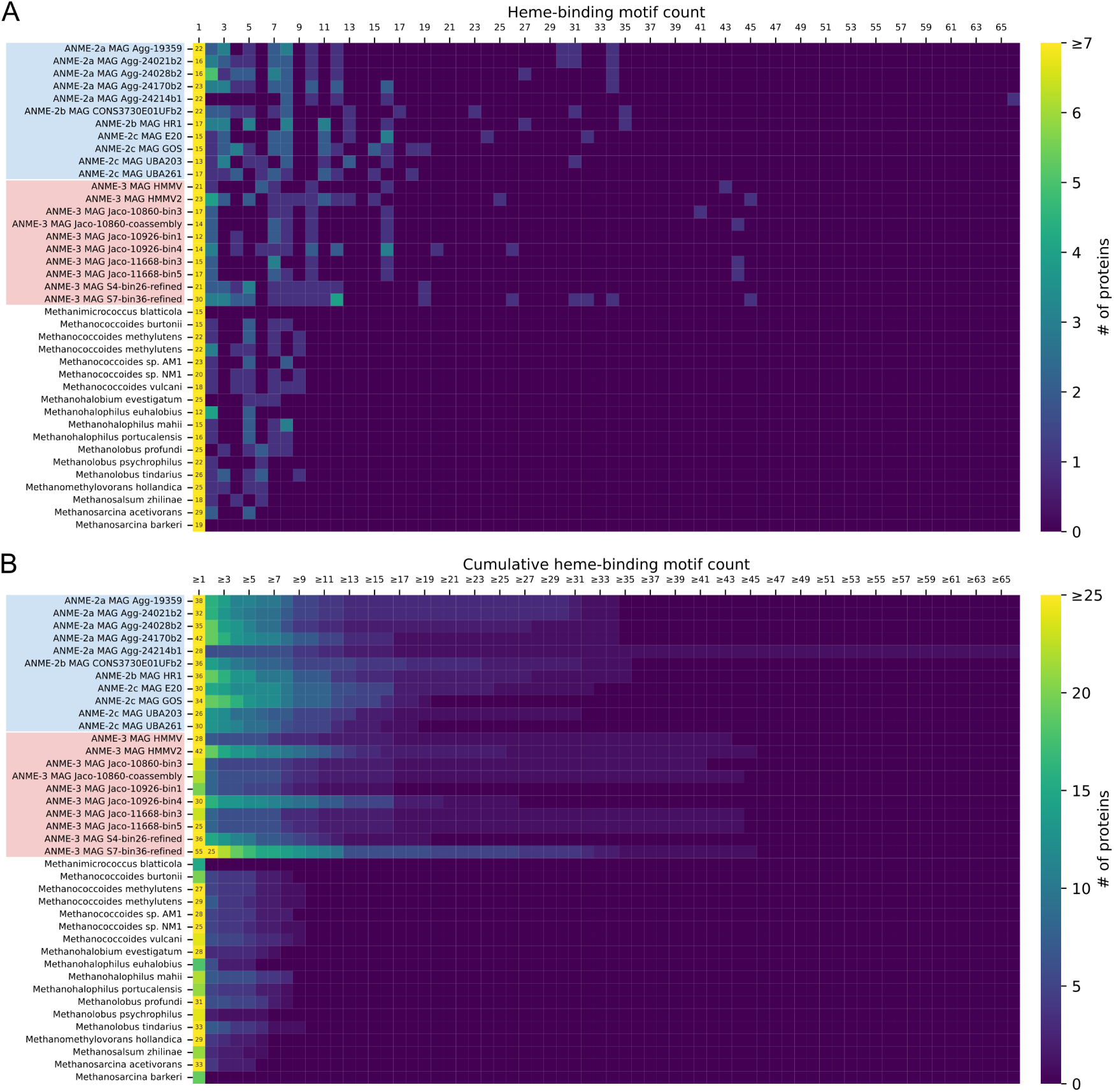
Distribution of putative cytochrome c proteins in ANME and methanogen genomes. (A) Heatmap showing the distribution of putative heme-binding motifs per protein in each genome. (B) Heatmap showing the cumulative distribution of putative heme-binding motifs per protein in each genome. All study genomes have similar numbers of single-heme proteins, but ANME have both more multi-heme cytochrome proteins overall and proteins with much higher numbers of heme binding motifs than related methanogens.

**Figure S10.**
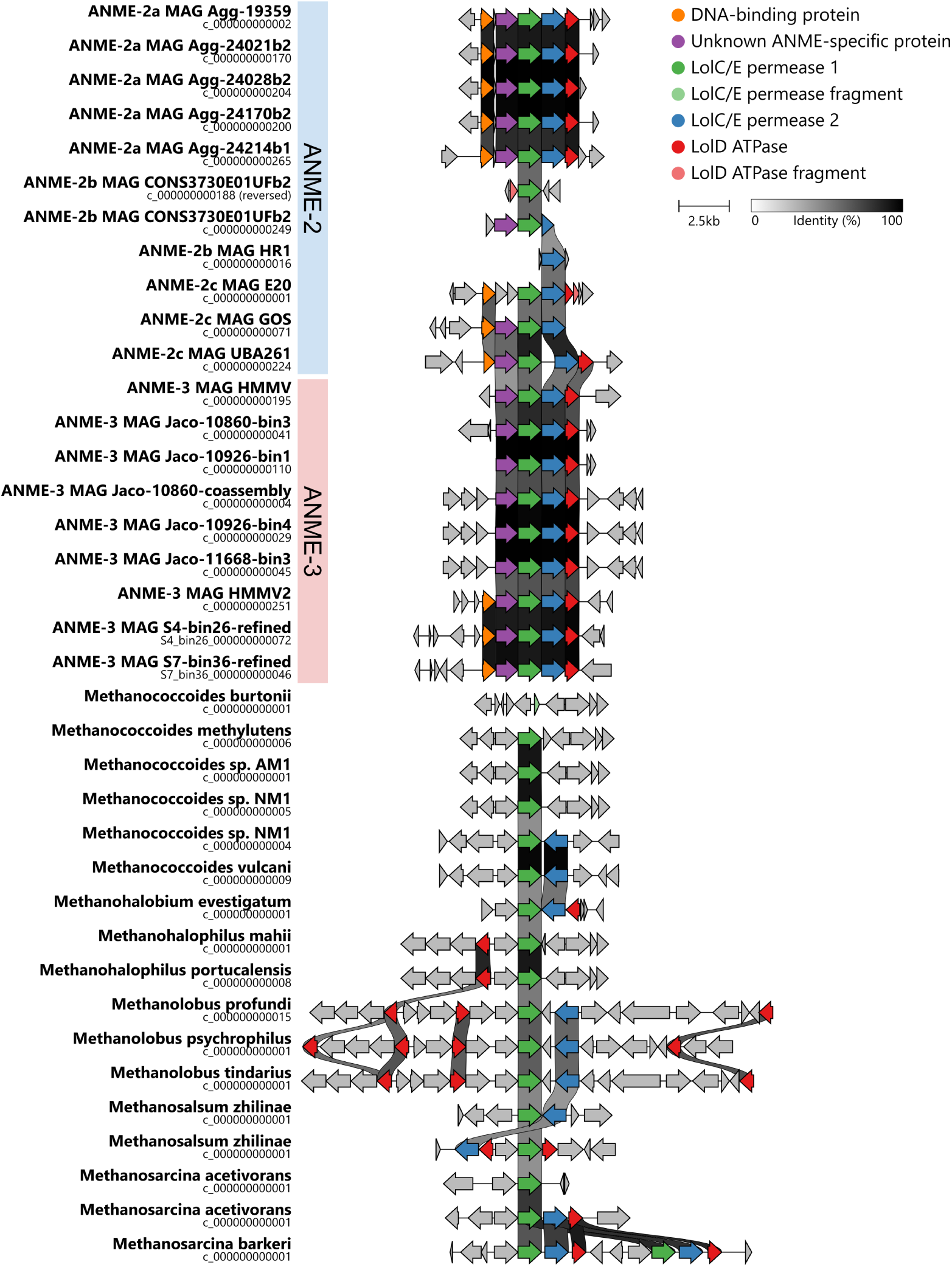
Synteny and gene content comparison of putative lipoprotein export locus. Proteins were grouped based on a 45% sequence identity threshold. ANME have highly consistent locus synteny and gene content, while methanogens vary in both organization and gene content. ANME loci also have moderately high sequence identity across genus and/or family boundaries.

**Figure S11.**
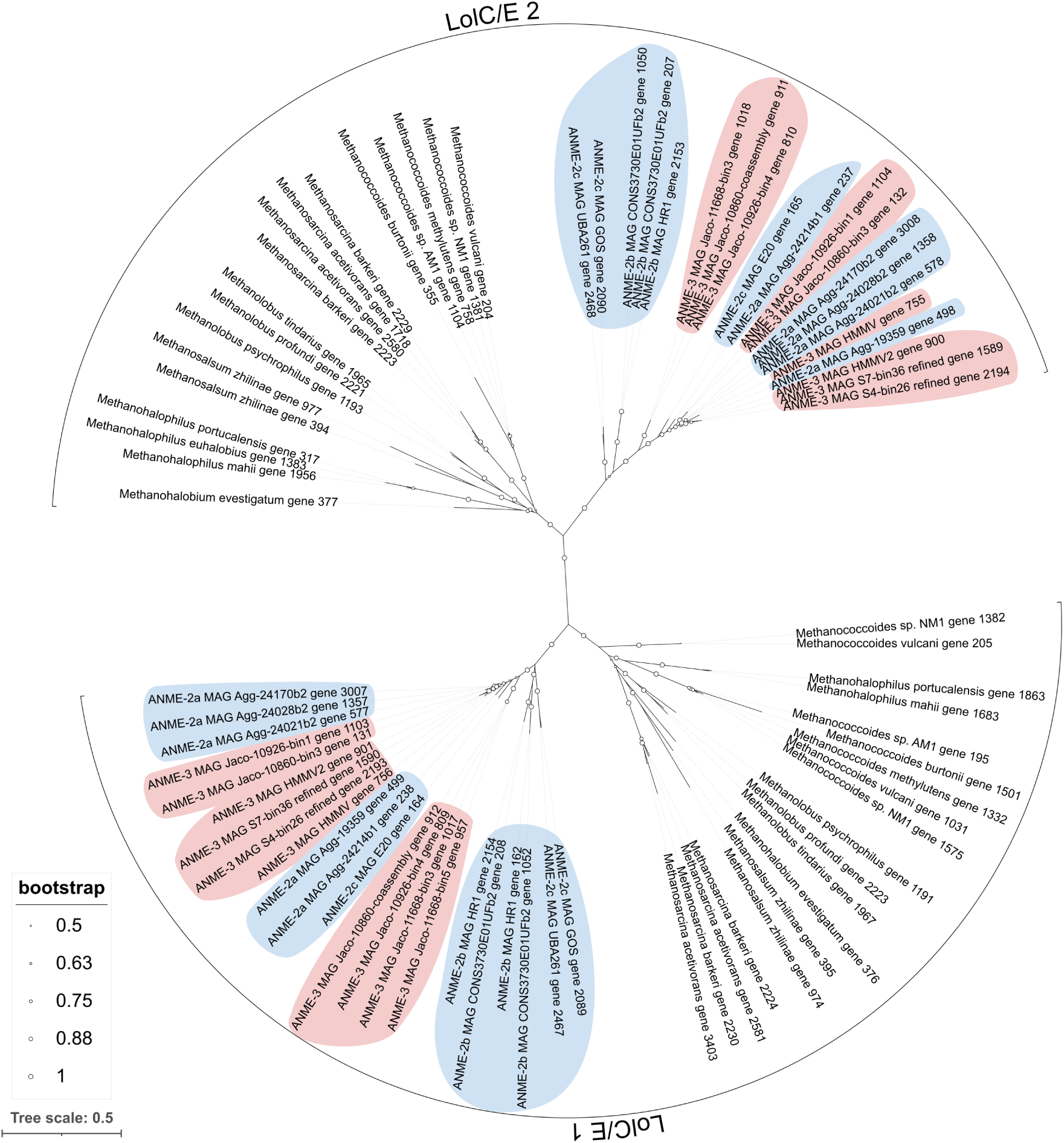
Gene phylogeny of putative lipoprotein export system genes *lolC/E* in ANME and methanogens. The two lolC/E genes appear to be homologous, and each gene shows clear separation between ANME and methanogen sequences. ANME-3 are interspersed among ANME-2 sequences for both genes. In some cases, ANME-3 sequences are grouped by inferred species phylogeny (e.g. the association between MAGs HMMV2, S4-bin26-refined, and S7-bin36-refined), but this is not always consistent (e.g. separation of Jaco MAGs).

**Figure S12.**
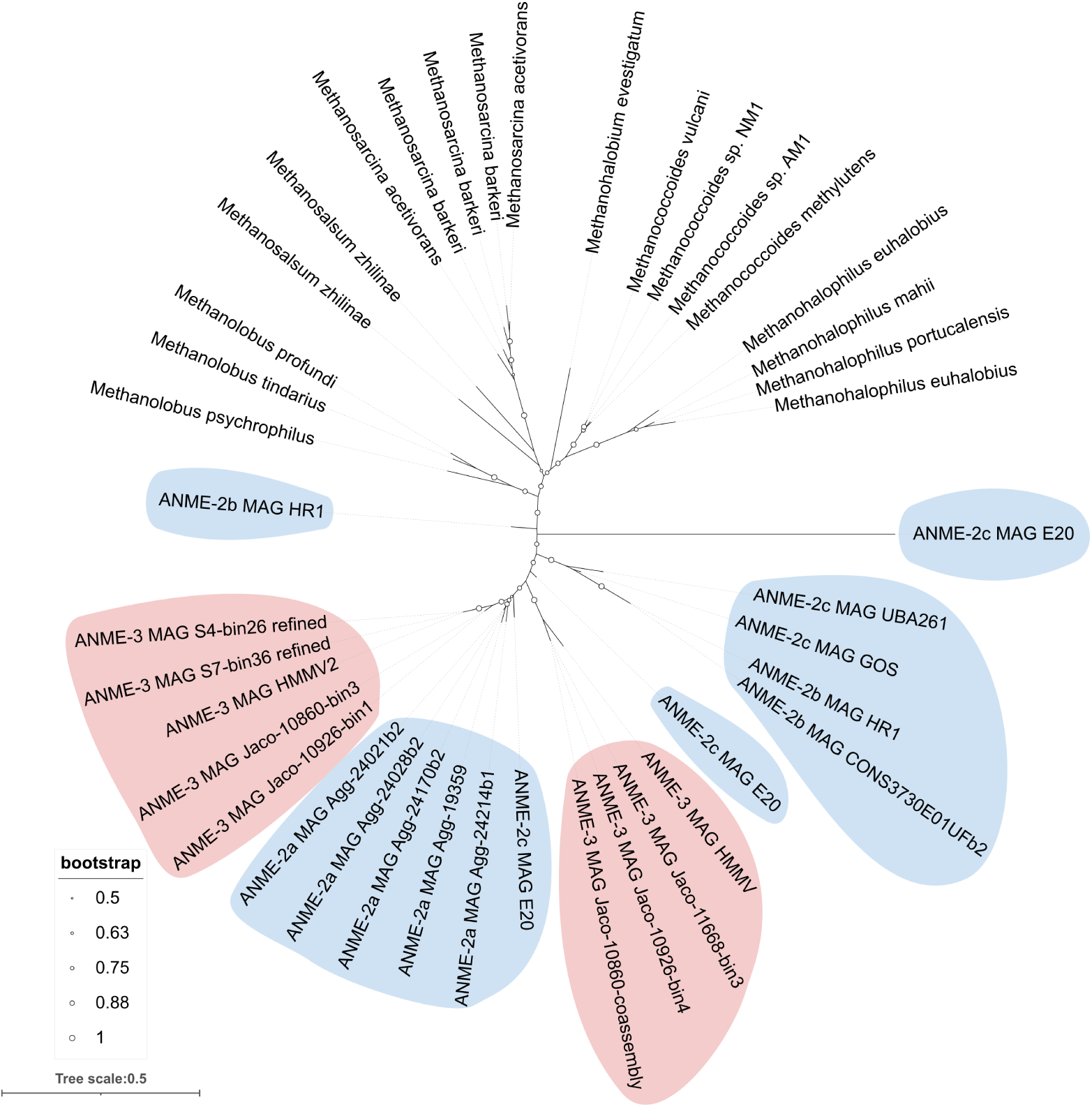
Gene phylogeny of putative lipoprotein export system gene *lolD* in ANME and methanogens. There is clear separation between ANME and methanogens. ANME-3 are interspersed among ANME-2 sequences. In some cases, ANME-3 sequences are grouped by inferred species phylogeny (e.g. the association between MAGs HMMV2, S4-bin26-refined, and S7-bin36-refined), but this is not always consistent (e.g. separation of Jaco MAGs).

**Figure S13.**
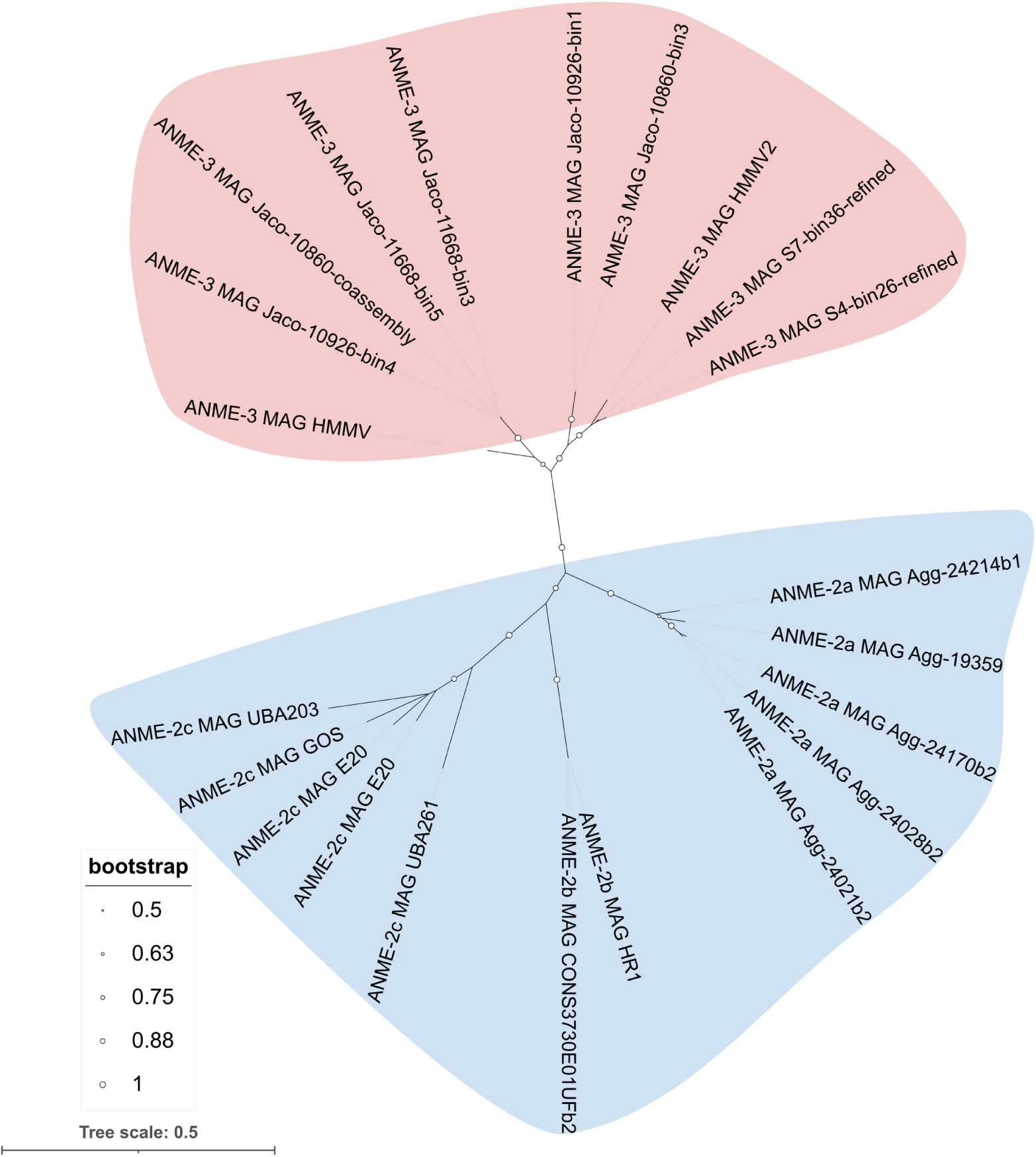
Gene phylogeny of unknown ANME-specific gene associated with putative lipoprotein export locus. Unlike other genes in this locus, sequences from ANME-2a, -2b, -2c, and ANME-3 are clearly separated into distinct clades. The gene phylogeny is similar to the estimated species phylogeny, though it does not perfectly agree for ANME-3 sequences.

**Figure S14.**
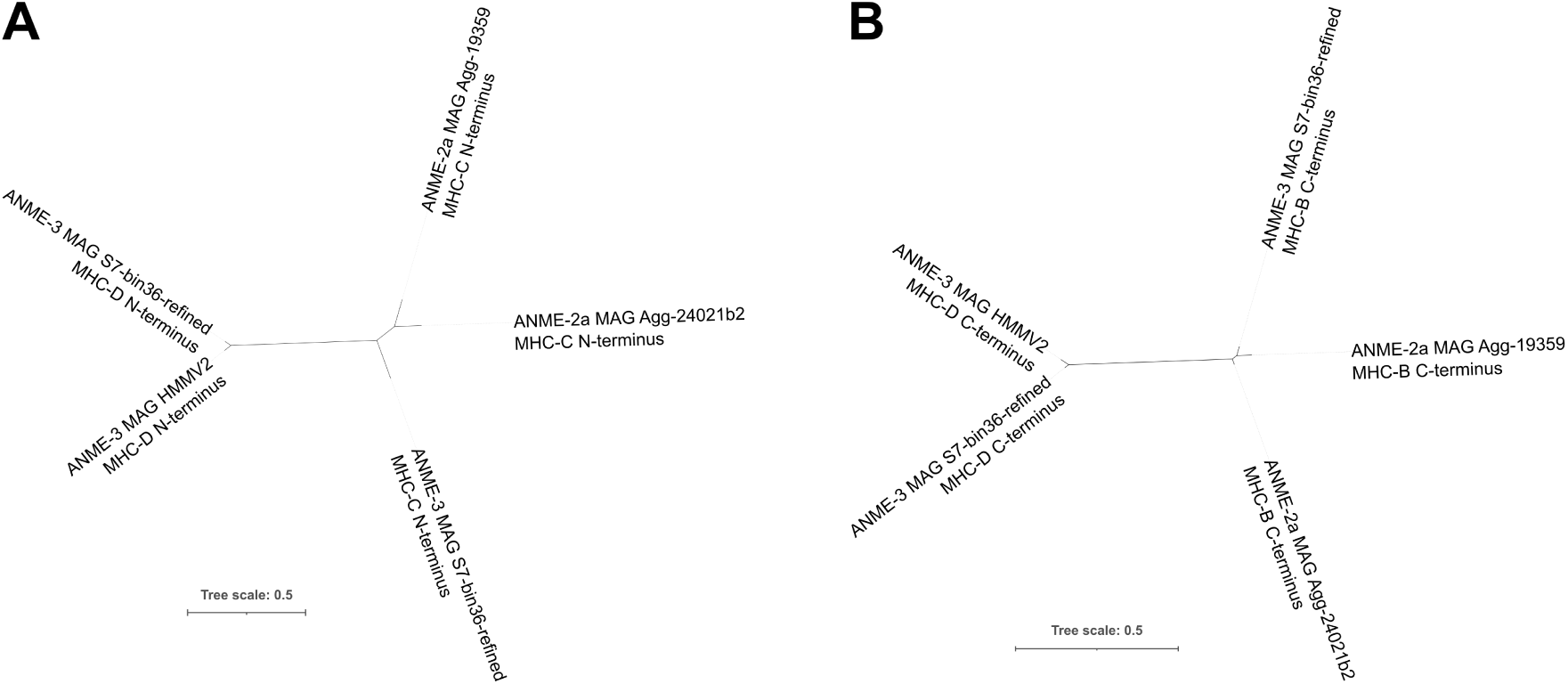
Gene phylogeny of the (**A**) N- and (**B**) C-terminal regions of MHC-D. The N-terminus of MHC-D aligns with the N-terminus of MHC-C, and the C-terminus of MHC-D aligns with the C-terminus of MHC-B. However, in both cases the MHC-D sequences cluster together to the exclusion of MHC-B or -C, even when both sequences are from the same genome as in the case in MAG S7-bin36-refined.

### 7.4 Supplemental files

S1. Function enrichments spreadsheet

S2. Pangenome summary spreadsheet

S3. RAxML settings

S4. Alignment of split PylS gene fragments to full-length sequences

S5. Full length McrA alignment

S6. Full length McrB alignment

S7. Full length McrG alignment

S8. McrA conservation report

S9. McrB conservation report

S10. McrG conservation report

S11. Full length MmcA alignment

S12. MHC-D N-terminus alignment

S13. MHC-D C-terminus alignment

S14. Core PLTS genes included and excluded from concatenated alignment

S15. MCR structure video

